# Insights into the structure and role of seed-borne bacteriome during maize germination

**DOI:** 10.1101/2020.06.02.130856

**Authors:** Lidiane Figueiredo dos Santos, Julie Fernandes Souta, Cleiton de Paula Soares, Letícia Oliveira da Rocha, Maria Luiza Carvalho Santos, Clicia Grativol Gaspar de Matos, Luiz Fernando Wurdig Roesch, Fabio Lopes Olivares

## Abstract

Seed germination events modulate microbial community composition, which ultimately influences seed to seedling growth performance. Here we assess the seed-borne bacteria community in disinfected and non-disinfected maize seeds and seedlings. Using a gnotobiotic system, sodium hypochlorite (1.25%, 30 min) treated-seeds showed a reduction of bacteria population size and an increase of bacteria community diversity associated with selective suppression of *Burkholderia* related taxon. The shift in the bacteria community composition in disinfested-seeds negatively affects germination speed, seedling growth, and reserve mobilization rates in comparison with non-disinfected maize seeds. A synthetic bacteria community formed by twelve isolates (9 *Burkholderia* spp.; 2 *Bacillus* spp. and 1 *Staphylococcus* sp.) obtained from natural microbiota of maize seeds herein were capable of recovering germination and seedling growth when reintroduced in disinfected seeds. Overall results showed that changes in bacterial community composition and selective reduction of *Burkholderia* related members dominance interfere with germination events and initial growth of the maize plantlets. By cultivation-dependent and independent approaches, we deciphered seed-maize microbiome structure, bacterial niches location, and bacterial taxon with relevant roles in seedlings growth performance. A causal relationship between seed microbial community succession and germination performance open opportunities in seed technologies to build-up microbial communities to boost plant growth and health.

**One sentence summary:** partial removal of the seed-borne microbiota negatively affects maize seedling growth performance and altered bacteria community structure. Partial microbial recomposition, mainly with *Burkholderia*-related isolates, restores the germination phenotype of disinfested seeds.

## INTRODUCTION

Soil microbial community represents the primary source of mutualistic microbes, which colonizes the soil rhizosphere of plants (Bakker, et al. 2015: 115-26, Hardoim, et al. 2012: e30438). Multi-omics approaches have revealed that soil harbor the most diverse bacteria population of any environment on earth (Fierer and Jackson 2006: 626-31, Roesch, et al. 2007: 283-90). From the soil continuum to the rhizosphere, plant-root exudates availability and composition are the major selective forces that shape rhizospheric microbiota, stimulating proliferation of certain bacterial specific groups (Bakker, et al. 2015: 115-26, Baudoin, et al. 2003: 1183-92). Presumably, such enriched bacterial populations have more chance to colonize root surface (epiphytically) and root inner tissue (endophytically), contributing to the significant fraction of the plant bacteriome composition.

Bacterial endophytes recruited from microbial diversity enriched by plant exudates have their soil-origin widely recognized (Bulgarelli, et al. 2012: 91-5). However, controversial reports have been published on the concern of the pivotal contribution of soil-borne bacteria for the plant bacteriome assembly (Johnston-Monje, et al. 2014: 233). Several studies pointed out the importance of seed-borne bacteria to build-up microbiome composition during plant growth and development (Nelson 2017: 7-34).

Vertically transmitted bacteria through seeds or vegetative plant parts have been extensively reported for different plant species. The underlying mechanisms of bacterial community dynamics and colonization of seeds, their offspring transmission, and predominant taxa under the germination process have been considered (Truyens, et al. 2015: 40-50).

More comprehensive studies about seed-associated microbes have been leveraged by cultivation-independent approaches (Adam, et al. 2016: 35-49, Hardoim, et al. 2012: e30438, Johnston-Monje, et al. 2016: 337-55). For instance, the 16S rRNA gene amplicon sequence allows broader assessment and comparison of the bacterial community of seeds, rhizosphere, and bulk soil. Thereby, the relative contribution of different factors (host genotypes, plant organ, and ontogeny, soil type, geography) that determine the assembly of seed endophytes can be adequately covered.

Among the plant species, maize seeds have received much attention as a niche for microbial communities (Truyens, et al. 2015: 40-50). Using culture-dependent techniques, (Rosenblueth, et al. 2010: 39-48) isolated distinct bacteria endophytes from surface-disinfected maize kernels detached from cobs and germinated under gnotobiotic conditions. In the early stages of germination, the predominant genera were *Bacillus* and *Paenibacillus*. Already *Methylobacterium*, *Alcaligenes*, *Tsukamurella*, *Erwinia*, *Microbacterium*, and *Rhodococcus* were detected later. Furthermore, *Burkholderia* was detected inside seeds using PCR of 16S rRNA. Bacterial endophytes were isolated from surface-sterilized maize kernels of four different cultivars under aseptic conditions, and the bacterial isolates were identified by 16S rRNA gene sequencing as *Pantoea* sp., *Microbacterium* sp., *Frigoribacterium* sp., *Bacillus* sp., *Paenibacillus* sp., and *Sphingomonas* sp. (Rijavec, et al. 2007: 802-8). Among eight bacteria isolated from the surface of disinfected seeds of thirty maize genotypes, *Bacillus* spp. was the most predominant species, with few isolates from the genera *Staphylococcus* and *Corynebacterium* (Bodhankar, et al. 2017: 232).

Cultivation-independent methods were applied to decipher the maize seed microbiome. Eight bacteria species were common in all six maize hybrids, representing a seed-inhabiting endophytic core. Among them, *Pantoea agglomerans*, *Enterobacter cloacae*, and *Aeribacillus palli* were the most representative taxa, accounting for 60% of relative abundance (Liu, et al. 2017: 317-24). The relative contribution from the soil and surrounding environment (horizontal transmission) and those inherited from seeds (vertical transmission) for endophytic assemblage in maize were accessed by (Johnston-Monje, et al. 2014: 233). It was concluded that the bacterial community from juvenile maize plants resembled a more seed bacteria community profile than the soil bacteria community in which the plants were growing. Also, around one-fifth of the bacterial community profiles harbored by the plant were typical in all soil geographical origin used, emphasizing the selective role of maize root exudates. In another study, it was demonstrated that despite the contribution of the soil bacteria for maize rhizosphere richness, the most dominant bacterial groups in juvenile maize rhizosphere are seed transmitted (Johnston-Monje, et al. 2016: 337-55).

We analyzed maize seed-borne bacteria by assessing the culture-dependent and culture-independent fraction of the bacterial community associated with germinated seedlings under gnotobiotic conditions. To gain insight into the role of epiphytic microorganisms, we used sodium hypochlorite as a disinfecting agent to eradicate mainly surface-attached microorganisms and then evaluated bacterial community structure, root-bacteria interaction by microscopy techniques and germination traits of maize seedlings. We have also assessed the effect of the reintroduction of a synthetic bacteria community formed by isolates obtained from the same maize seed-borne genotype herein.

## MATERIAL AND METHODS

### Disinfection and germination of maize seeds

Seeds of commercial maize (*Zea mays* L.) hybrid SHS 5050 (Santa Helena Sementes, Brazil) were washed five times in sterile distilled water, remaining for 5h in the last water. After the imbibition period, the seeds were divided into two treatments: a) Non-disinfected seeds (NDS) and b) Disinfected seeds (DS). For DS-treatment, seeds were surface disinfected in 70% ethanol for 5 min and soaked in sodium hypochlorite (NaClO; Butterfly Ecologia, Audax Company) at 1.25% for 30 min with subsequent five rinses in sterile distilled water. The experimental unit comprised twelve seeds from NDS or DS treatment aseptically placed in a Petri dish containing agar-water medium (0.5%) under axenic conditions. Five replicates were used for each treatment in a completely randomized design (DIC). Non-disinfected and disinfected seeds were germinated in a growth chamber at 30 °C and photoperiod 12/12 h (light/dark) for 5 and 8 days, respectively. During this period, the number of germinated seeds (radicle ≥ 5 mm) was recorded daily to calculate different germination parameters, including germination percentage (%G), germination speed index (GSI), average germination time (AGT) and average germination speed (AGS) (Maguire 1962: 176-7), 1962). Significance between treatments was calculated using an unpaired t-test (p ≤ 0.05). The disinfection time was defined by experimentally (Supplementary Figure S1).

### Estimation of bacteria population associated with germinated seeds and isolation of maize seed-borne bacteria

The population size of total and diazotrophic bacteria were estimated with root samples obtained in the germination assay for disinfected and non-disinfected maize seeds (Table 1). For this, 1 g of root from each Petri dish was macerated in saline 99 mL of sodium chloride (NaCl, 8.5 g L^−1^) and subjected to serial dilution from 10^−3^ to 10^−6^. After that, 100 μL of each dilution was transferred to culture media. Plates were incubated in the growth chamber at 30 °C for 5 to 7 days. Solid NB medium was used for counting total bacteria, and results were expressed in colony-forming units per g (fresh weight root). For diazotrophic bacteria count, two media were used, the JNFb semi-solid medium (malic acid as C-source) and LGI semi-solid medium (sucrose as C-source). The media composition and procedure to determine Most Probable Number (MPN) per g of a fresh root followed the recommendation of (Baldani, et al. 2014: 413-31).

**Table 1.**
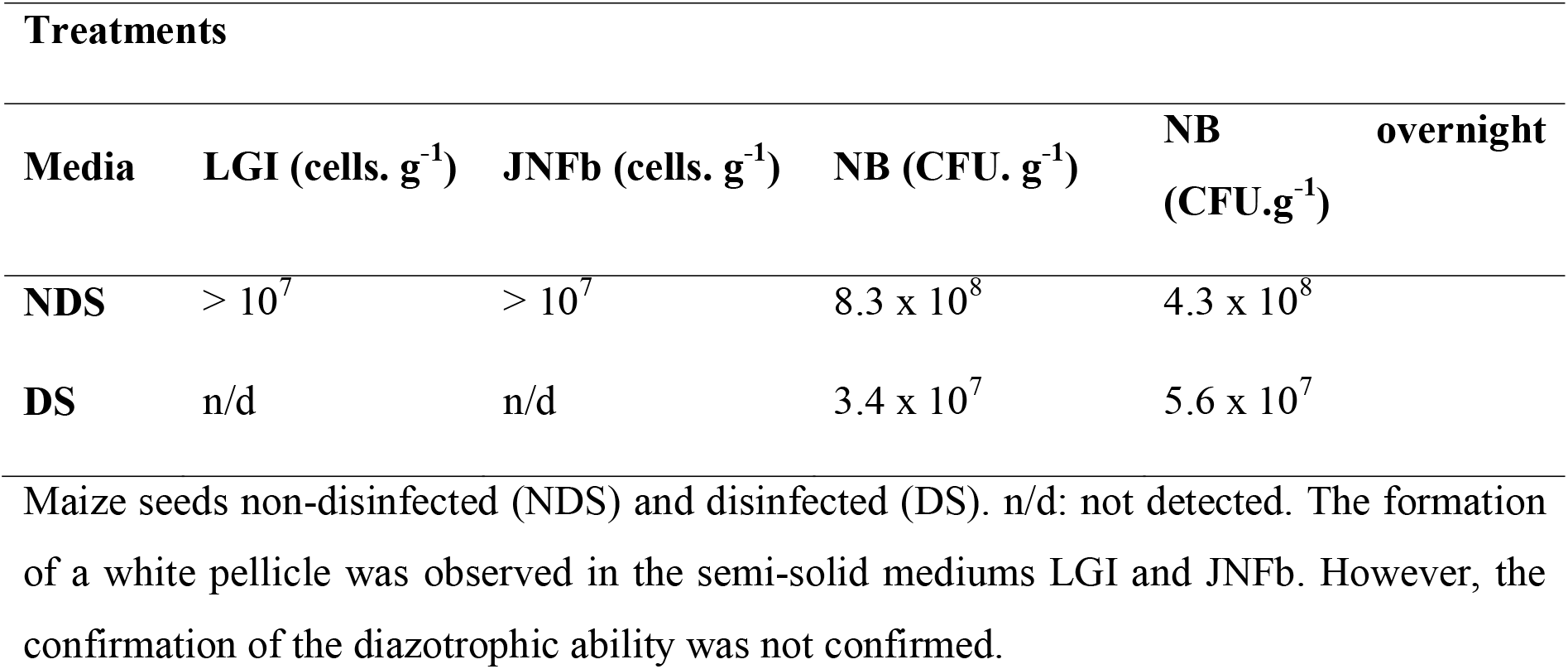
Count of total and diazotrophic bacteria recovered from both NB solid media and LGI and JNFb semi-solid media.

Additionally, whole primary root maize segments were soaked in 5 mL tubes with liquid NB medium, kept under agitation overnight (180 rpm, at 30 °C). Then, the liquid medium was subjected to serial dilution and plated in solid NB, for colony counting. Bacteria isolates obtained from different sources and media were checked for purity in NB solid medium. Purified colonies and cell morphology were described, according to (Baldani, et al. 2014: 413-31). Colony diversity analysis was performed by calculating Shannon’s abundance and index in the Past program, version 3 (Supplementary Figure S2).

### Microscopy evaluation of the maize seed microbiota

Non-germinated (Supplementary Figure S3) and germinated maize seeds (Figure 1–2) of disinfected and non-disinfected seeds were collected and processed for LM and SEM aiming to characterize the seed and root-associated bacterial microbiome.

**Fig 1.**
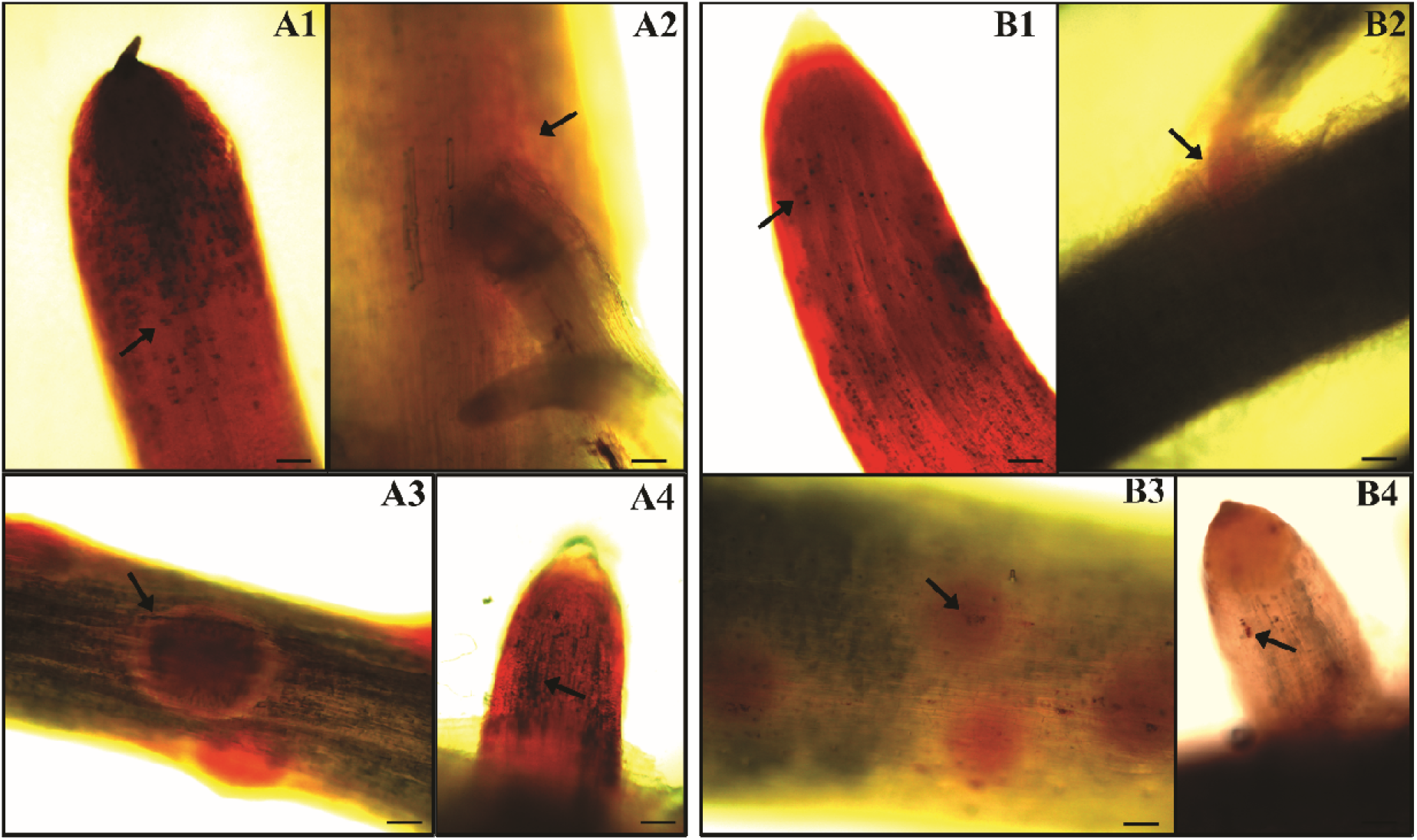
Bacterial colonization of maize roots stained with triphenyl tetrazolium chloride (TTC). The cells were visualized by light microscopy. Maize seeds non-disinfected (A) and disinfected (B). Root regions: root cap (A1; B1), lateral root emergence (A2 and B2; A4 and B4), and mitotic sites (A3; B3). Black arrows indicate small bacterial aggregates. Bars represent the following scales 10 μm.

**Fig 2.**
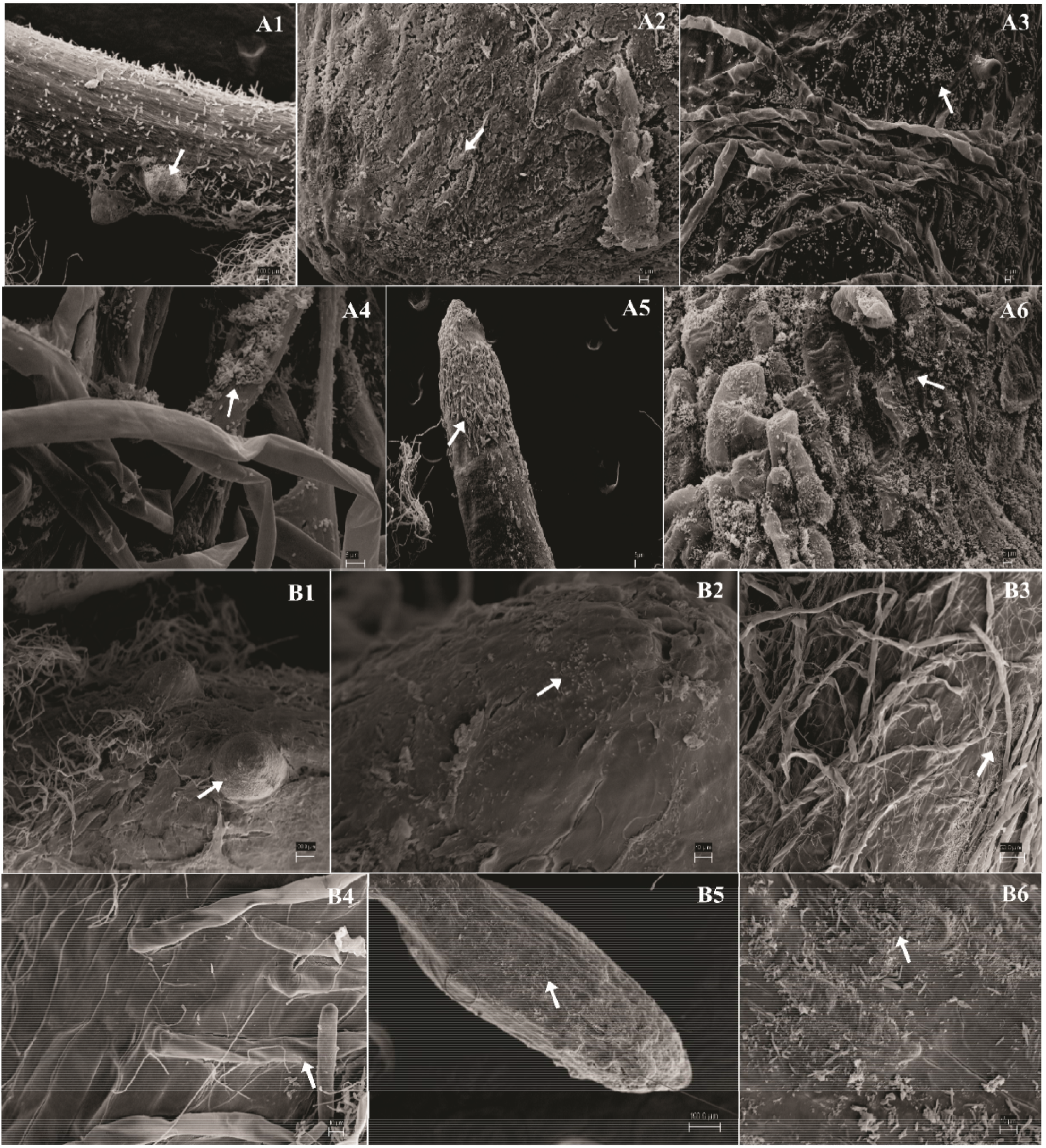
Colonization of maize roots by the microbiota. Bacterial cells were visualized by scanning electron microscopy (SEM). Maize seeds non-disinfected (A) and disinfected treatments (B). Root regions: mitotic sites (A1 and A2; B1 and B2), by the roots (A3 and A4; B3 and B4) and cap (A5 and A6; B5 and B6). White arrows indicate biofilms and small bacterial aggregates. Bars represent the following scales: panel A2, A3, A4, A5, and A6: 5 μm; B2, B4, and B6: 10 μm; B3: 50 μm; A1, B1, and B5: 100 μm.

From the germination assay described herein, the whole primary root of the maize seedlings was obtained from each treatment. The fresh roots segment was immersed and stained with 2,3,5-triphenyl tetrazolium chloride (0.1% TTC for 2h) and then dipped in potassium hydroxide solution (2.5% KOH for 40 min) to reduce the staining background of the root tissue. The colored material was placed on glass slides, mounted with sterile distilled water, and visualized under bright-field microscope Axioplan Zeiss equipped with AxioCam digital photography system (Figure 1).

For SEM, root segments (≈ 1 cm) and transversal/longitudinal sections of whole seeds were fixed in a solution containing 2.5% glutaraldehyde and 4% paraformaldehyde in potassium phosphate buffer (0.05 mol L^−1^, pH 7.0). Then, the samples were washed (3 times for 20 min for root; 30 min for seed) in the same buffer and dehydrated in a growing series of ethanol (15, 30, 50, 70, 90 and 2 × 100% at 15 min for each root; 30 min for seed). After dehydration, samples were dried in critical point drier apparatus (BAL-TEC CPD 030), mounted on Al-stubs, and metalized with ionized platinum in a sputtering coat apparatus (BAL-TEC SCD 050); and visualized under an SEM Zeiss EVO 40 (Figure 2). Macroscopic structures of the seeds were recorded using a magnifying glass (Zeiss Stemi SV 11) coupled to a digital camera and used as a reference for the scanning images.

### Sequencing of the bacterial microbiome

Samples of disinfected and non-disinfected non-germinated seeds (Supplementary Figure S4-S5) and germinated seeds and roots (Figure 3–4) were macerated in liquid nitrogen and stored at −70 °C until the extraction of the genetic material. The total genomic DNA was isolated from seeds and roots (~0.2 g) using two adapted extraction protocols: 1) Plant DNAzol® kit, for roots (Ausubel, et al. 1990, Wilfinger, et al. 1997: 474-81); 2) Cetyltrimethylammonium bromide (CTAB), for roots and seeds (Chen and Ronald 1999: 53-7, Doyle and Doyle 1990: 13-5). Then, the DNA was quantified by a NanoDrop 2000® spectrophotometer (Thermo Scientific) and Qubit® fluorometer (Invitrogen); and its quality evaluated on agarose gel (0.8%) electrophoresis (80 V, for 70 min). The total DNA was sent to the company “WEMSeq Biotecnologia” for sequencing of the 16S rRNA gene in Illumina MiSeq, with three replicates per treatment. Twenty nanograms of DNA were used as a model for 18 cycles of amplification of the V4 region of the 16S rRNA gene, using primers 515F and 806R (Caporaso, et al. 2012: 1621-4) and GoTaq Master Mix (Promega). PCR products were quantified with the Qschd dsDNA HS kit (Invitrogen) and sequenced with the 300V2 Kit (Illumina) in Illumina MiSeq (Illumina), following the manufacturer’s instructions.

**Fig 3.**
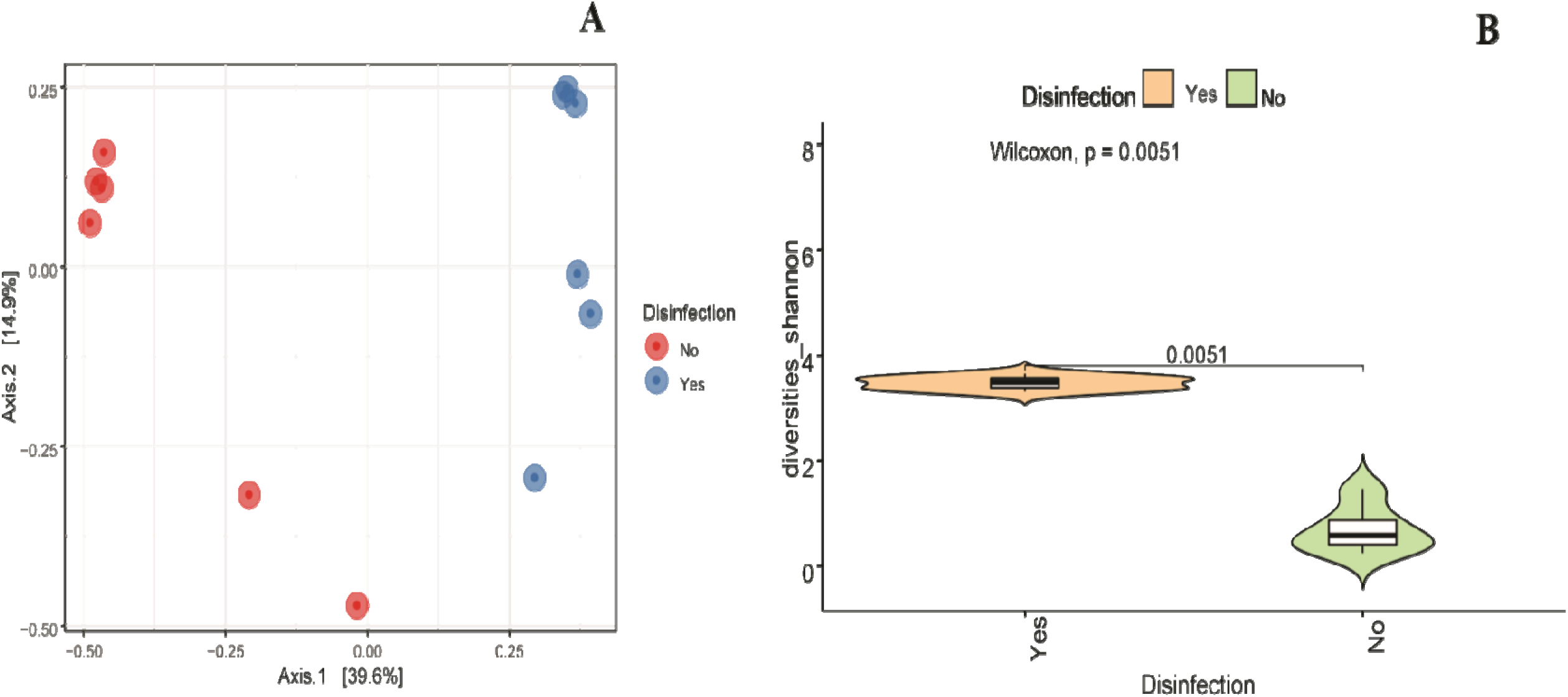
The principal coordinate graph (PCoA) based on the Bray-Curtis dissimilarity matrix (A) and measurements of the alpha diversity of root bacteriome from disinfected (Yes) and non-disinfected (No) seeds. Different colors indicate the treatments. Shannon = microbial diversity index.

**Fig 4.**
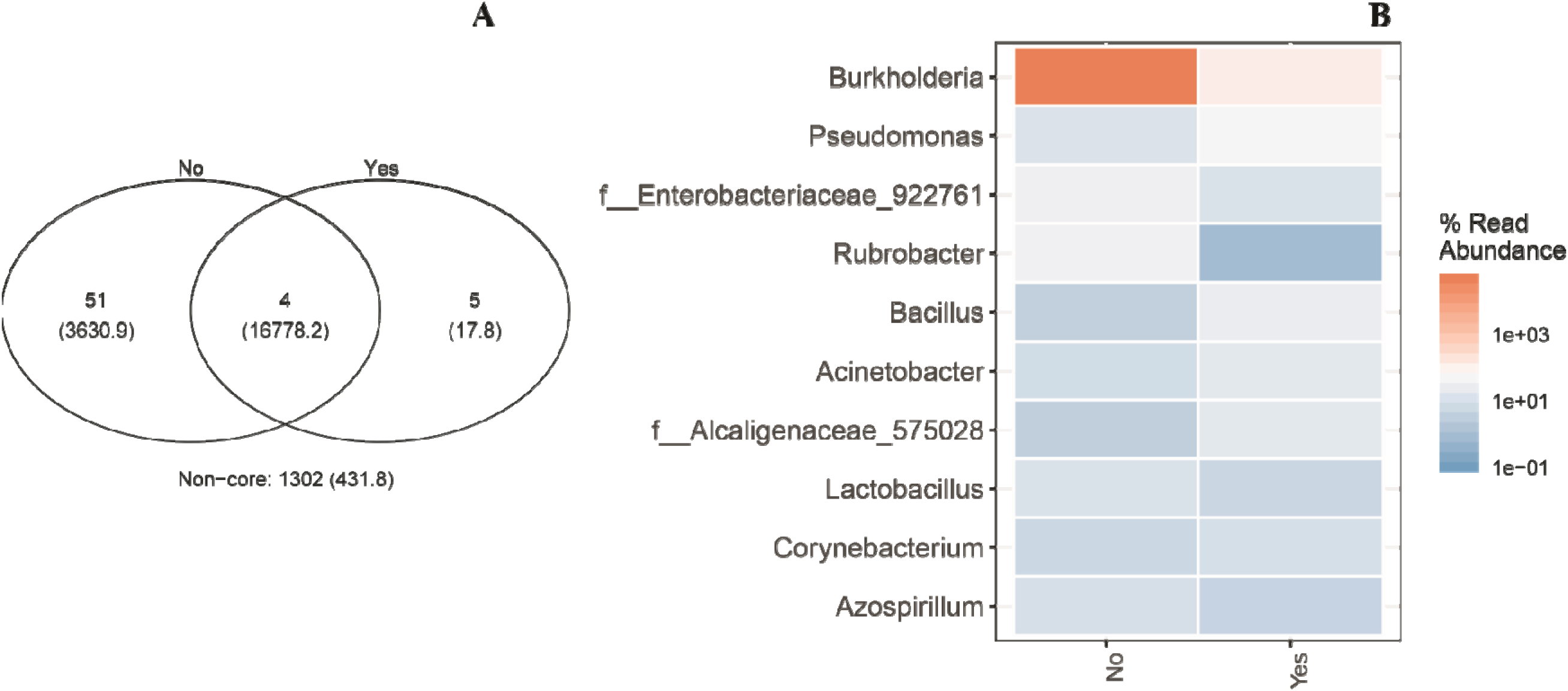
Venn diagram showing OTUs overlap (A) and the relative abundance between roots from disinfected (Yes) and non-disinfected (No) seeds (B). The color intensity of heatmap indicated in the legend to the right of the figure shows the relative values for the genus.

The sequences obtained from MiSeq were analyzed using the QIIME (Quantitative Insights Into Microbial Ecology) version 1.9.0 (Caporaso, et al. 2010: 335), where they were filtered and grouped into Operational Taxonomic Units (OTUs) in five levels: phylum, class, order, family, and genus, considering 97% similarity with the SILVA database (Quast, et al. 2012: D590-D6). BIOM files were imported into the R environment using the phyloseq package (McMurdie and Holmes 2013). Sampling quality was estimated using Good’s coverage. To test the hypothesis that disinfection shapes the bacteriome, principal coordinate analysis (PCoA) graphs were constructed based on the Bray-Curtis dissimilarity matrix. Differences between treatments were obtained by Multivariate Analysis of Permutational Variance (PERMANOVA) (Anderson 2014: 1-15). Alpha diversity was estimated by applying the Shannon diversity index, in addition to calculating the differential abundance for root samples. Venn diagrams and genus-level heatmaps were built.

### Quantitative PCR (qPCR) for eubacteria

Disinfected and non-disinfected maize seeds and roots had their bacterial community quantified by qPCR from DNA extracted (Figure 5) according to the methodology mentioned above (chosen method: CTAB). Each PCR reaction (15 μL) contained template DNA (100 ng for seed and 40 ng for root); 7.5 μL of SYBR Green (Promega), 0.5 μL of each primer (10 μM; 926F: AAACTCAAAKGAATTGACGG; 1062R: CTCACRRCACGAGCTGAC) (De Gregoris, et al. 2011: 351-6) and water. The reactions were carried out in triplicate, with 5 min incubation at 95 °C, 40 cycles of 15 s at 95 °C and 1 min at 60 °C in Step-One-Plus Real-Time PCR System (Applied Biosystems). The proportion of bacteria in the microbiome was calculated based on the Ct (cycle threshold) values of the samples (three replicates per treatment) and using a standard curve generated from the model bacteria *Escherichia coli* ATCC 25922. *E. coli* was grown in NB liquid medium (180 rpm, at 30 °C) and had its DNA extracted with Wizard Genomic DNA Purification Kit (Promega). The reference DNA was diluted in series of 10^−2^ to 10^−8^ (20 – 2 × 10^9^ ng of DNA), and its quantification in qPCR (expressed in Ct) plotted concerning the number of *E. coli* cells.

**Fig 5.**
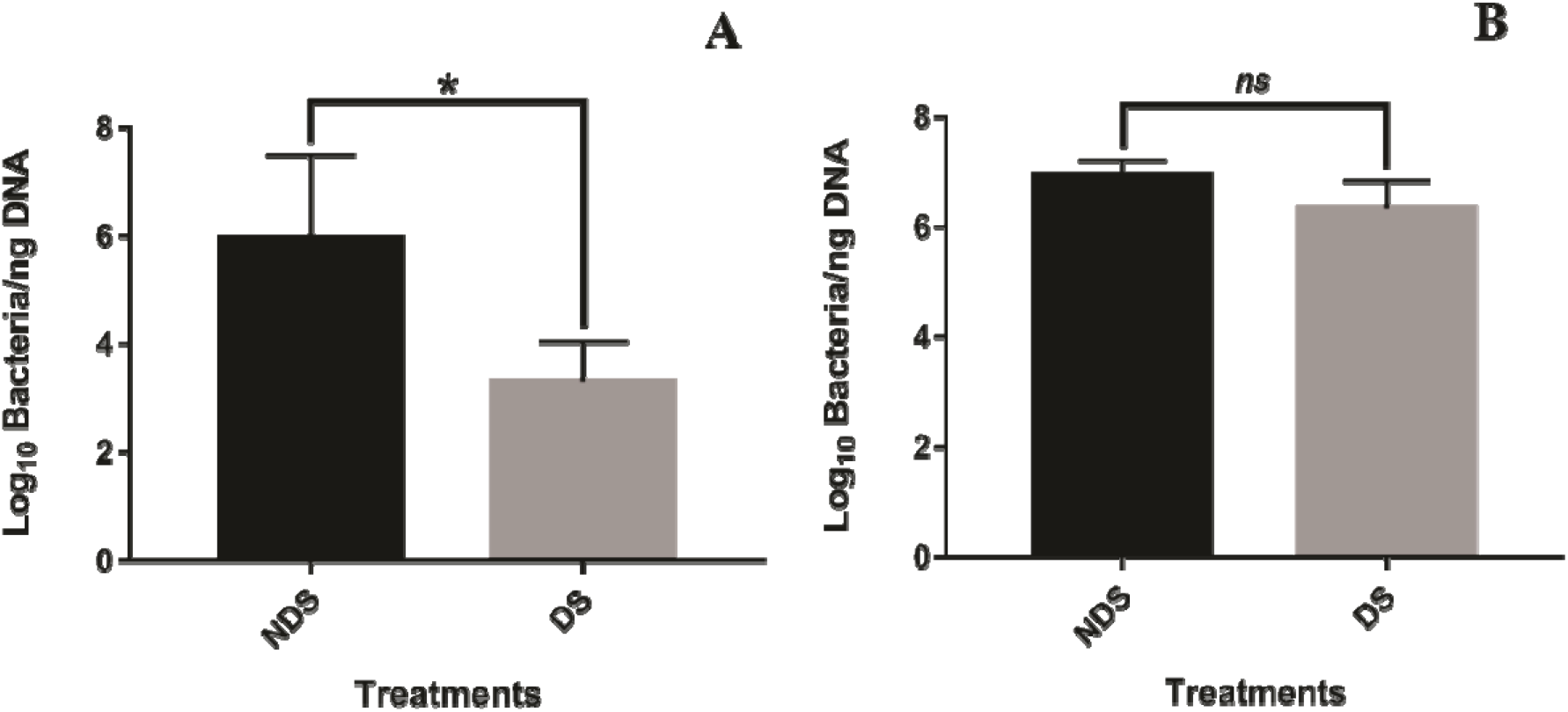
Quantification of microbiome bacteria by qPCR in non-disinfected (NDS) and disinfected (DS) seeds (A) and roots (B). *Significant difference between treatments according to the Tukey test (p ≤ 0.05).

### Partial recomposition of the bacterial maize microbiota

Bacteria isolates obtained from non-disinfected maize seeds axenically were recovered from NB solid medium and JNFb or LGI semi-solid medium herein. Based on colony morphology diversity, twelve bacteria isolates were selected for partial recomposition microbiota assays. Strain designation and origin of the isolates are quoted in Supplementary Table S1. The taxonomic affiliation of the selected bacterial isolates can be seen in Supplementary Table S2. Each bacterium of this consortium was grown in liquid NB medium, kept under agitation (180 rpm, for 24-48 h at 30 °C). The bacterial cells were collected by centrifugation (5,000 rpm, for 10 min (Eppendorf)) and resuspended in sterile distilled water. The values of OD595 (optical density at 595 nm) of the isolates were standardized on a spectrophotometer (OD595 = ~0.5 in NB medium), and these were mixed to form a synthetic bacteria community diluted until 10^−7^ concentration from an initial cell density of 2 × 10^8^ UFC.g-1.

For the inoculation of the synthetic bacteria community, disinfected maize seeds were immersed in the inoculum for 10 min, aiming at the partial rebuilding of the microbiota removed by seed disinfection with sodium hypochlorite. Then, the seeds that were disinfected and that received the mix of 12 bacteria (DS + MIX) were placed in Petri dishes with agar-water medium (0.5%). Ten seeds per plate constituted the experimental plot. The experimental design was completely randomized with the following treatments: 1) NDS; 2) DS; 3) DS + MIX and five repetitions for germination test evaluation and four repetitions for the plant-growth evaluation (Fig 6–7). The disinfected and non-disinfected seeds were used as controls. All plates were conditioned in the growth chamber at 30 °C and photoperiod 12/12 h (light/dark) for five days. Germination rate was calculated, and the length and mass (fresh and dry) of the aerial part and root seedlings with the aid of a millimeter ruler and analytical balance. The counting of total bacteria and the visualization of cells in maize roots stained with TTC by light microscopy were also carried out, according to methodologies previously described (Figure 9). Scanning electron microscopy was used for the structural characterization of reinoculated seeds and roots, respectively (Figure 8–10). The results were submitted to analysis of variance (ANOVA) and the means compared by the Tukey test (p ≤ 0.05).

**Fig 6.**
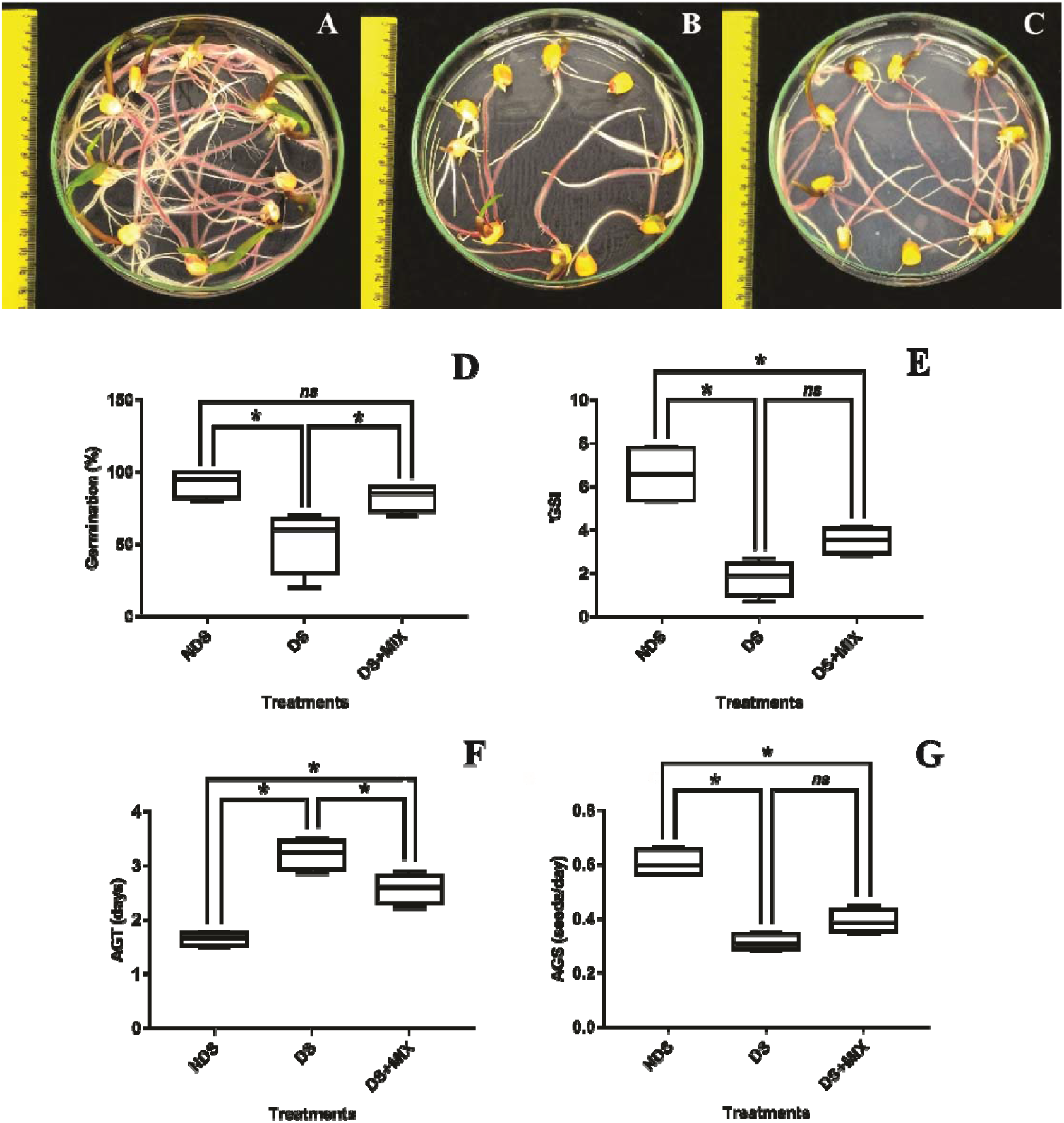
Maize seeds non-disinfected (A; NDS), disinfected (B; DS), and disinfected-inoculated (C; DS + MIX). Germination percentage (D), germination speed index (E), average germination time (F), and average germination speed (G). *Significant difference between treatments according to the Tukey test (p ≤ 0.05).

**Fig 7.**
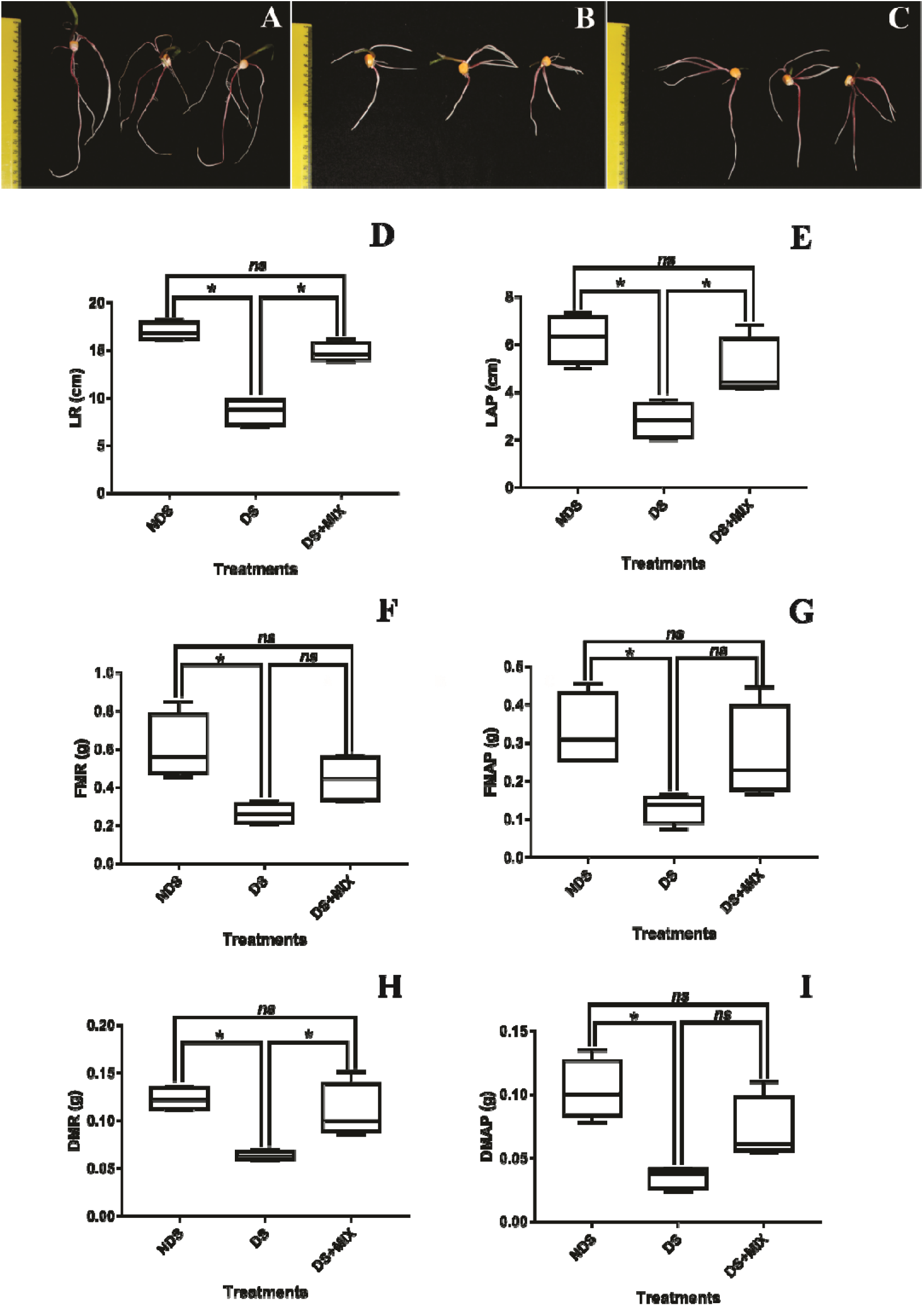
Maize seeds non-disinfected (A; NDS), disinfected (B; DS), and disinfected-inoculated (C; DS + MIX). Shoot length (D) and root (E), shoot fresh weight (F) and root (G), shoot dry weight (H), and root (I). *Significant difference between treatments according to the Tukey test (p ≤ 0.05).

**Fig 8.**
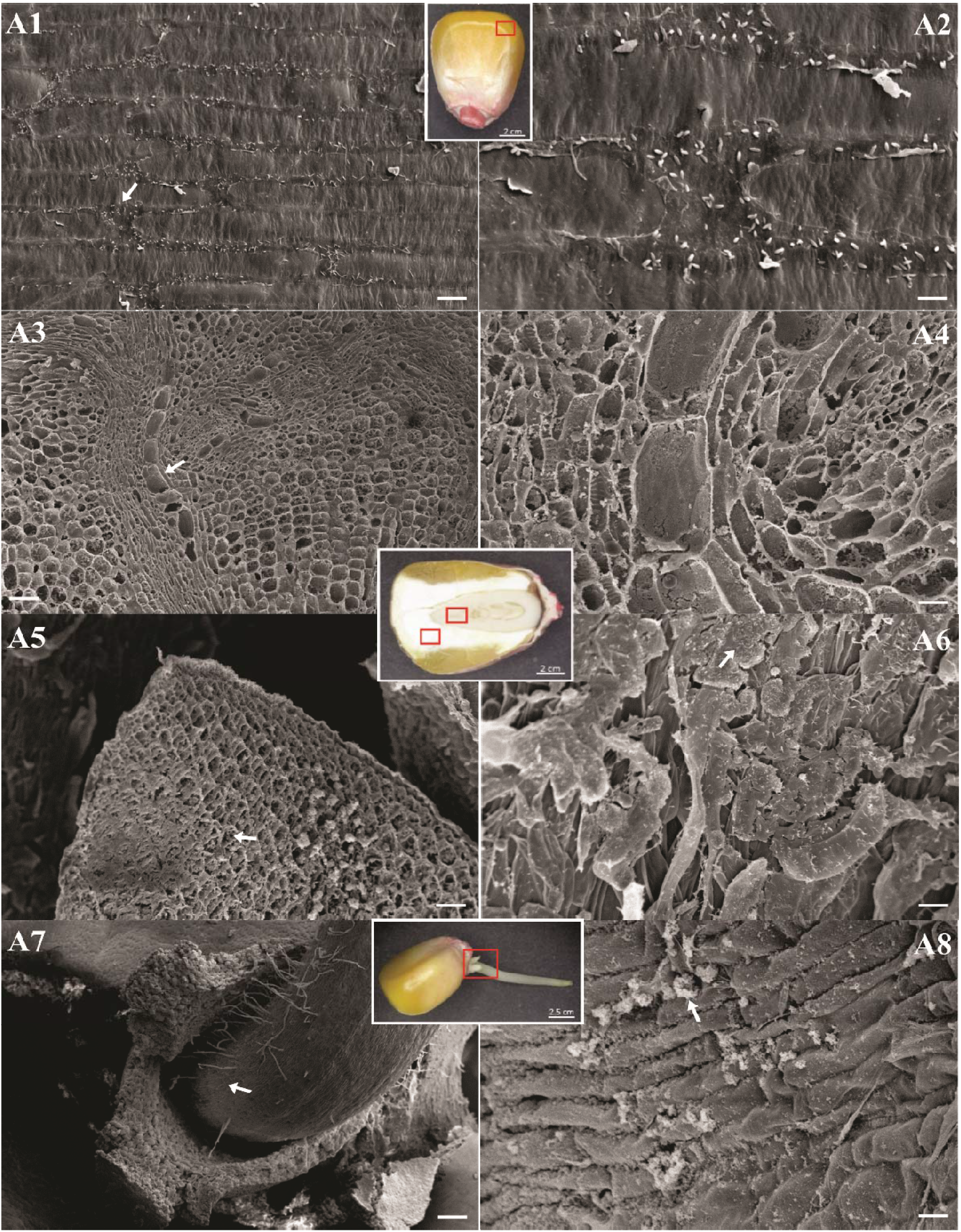
Colonization of disinfected-inoculated maize seeds (A). Bacterial cells were visualized by scanning electron microscopy. Seed regions: pericarp (A1 and A2), endosperm (A3 and A4), embryo (A5 and A6), and radicle (A7 and A8). White arrows indicate biofilms and small bacterial aggregates. Bars represent the following scales: panel A2, A6, and A8: 10 μm; A1 and A4: 20 μm; A3 and A5: 100 μm; A7: 200 μm.

**Fig 9.**
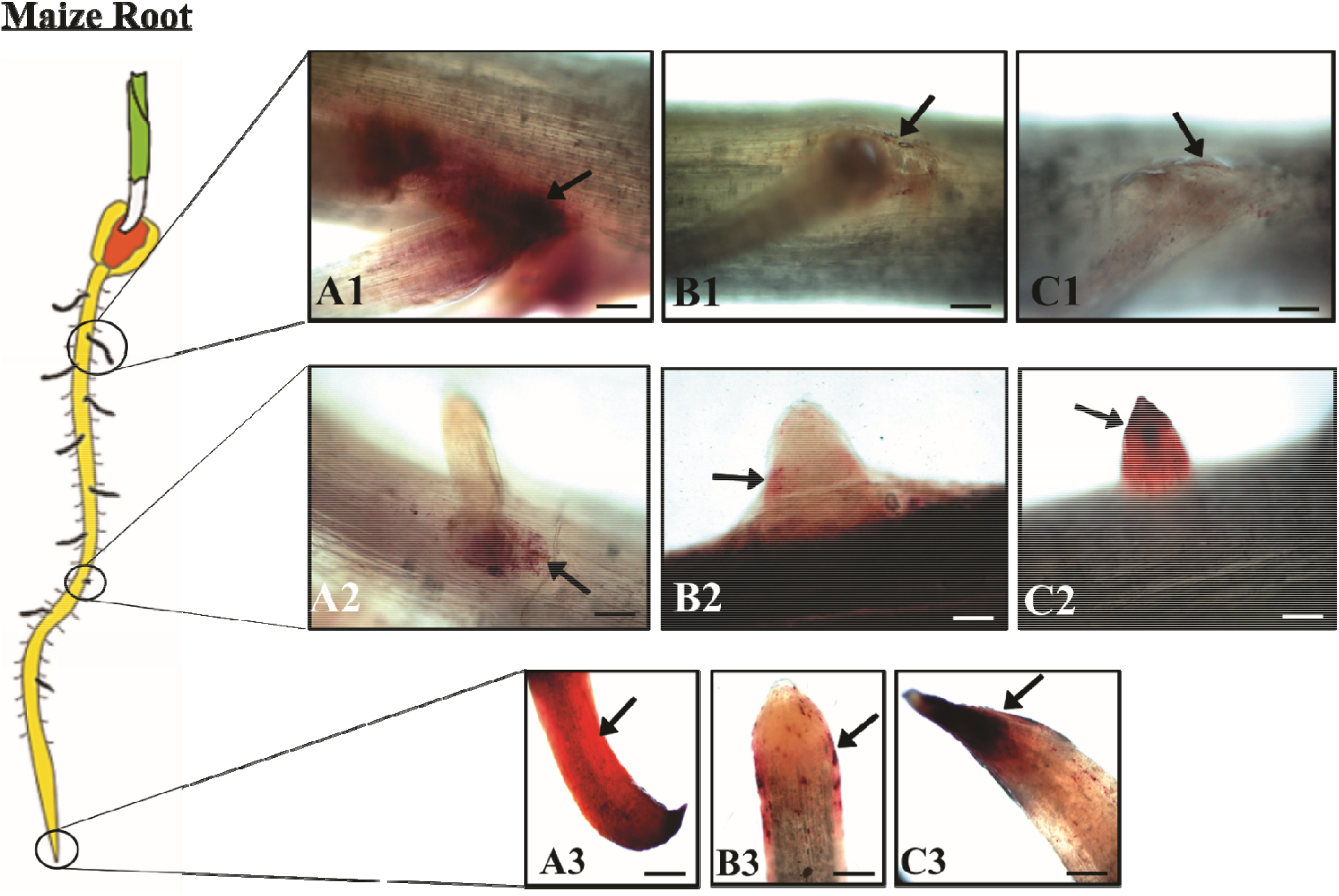
Bacterial colonization of maize roots stained with triphenyl tetrazolium chloride (TTC). The cells were visualized by light microscopy. Maize seeds non-disinfected (A), disinfected (B), and disinfected-inoculated (C). Root regions: lateral root (A1 and A2; B1 and B2; C1 and C2) and cap (A3; B3; C3). Black arrows indicate small bacterial aggregates. Bars represent the following scales 10 μm.

**Fig 10.**
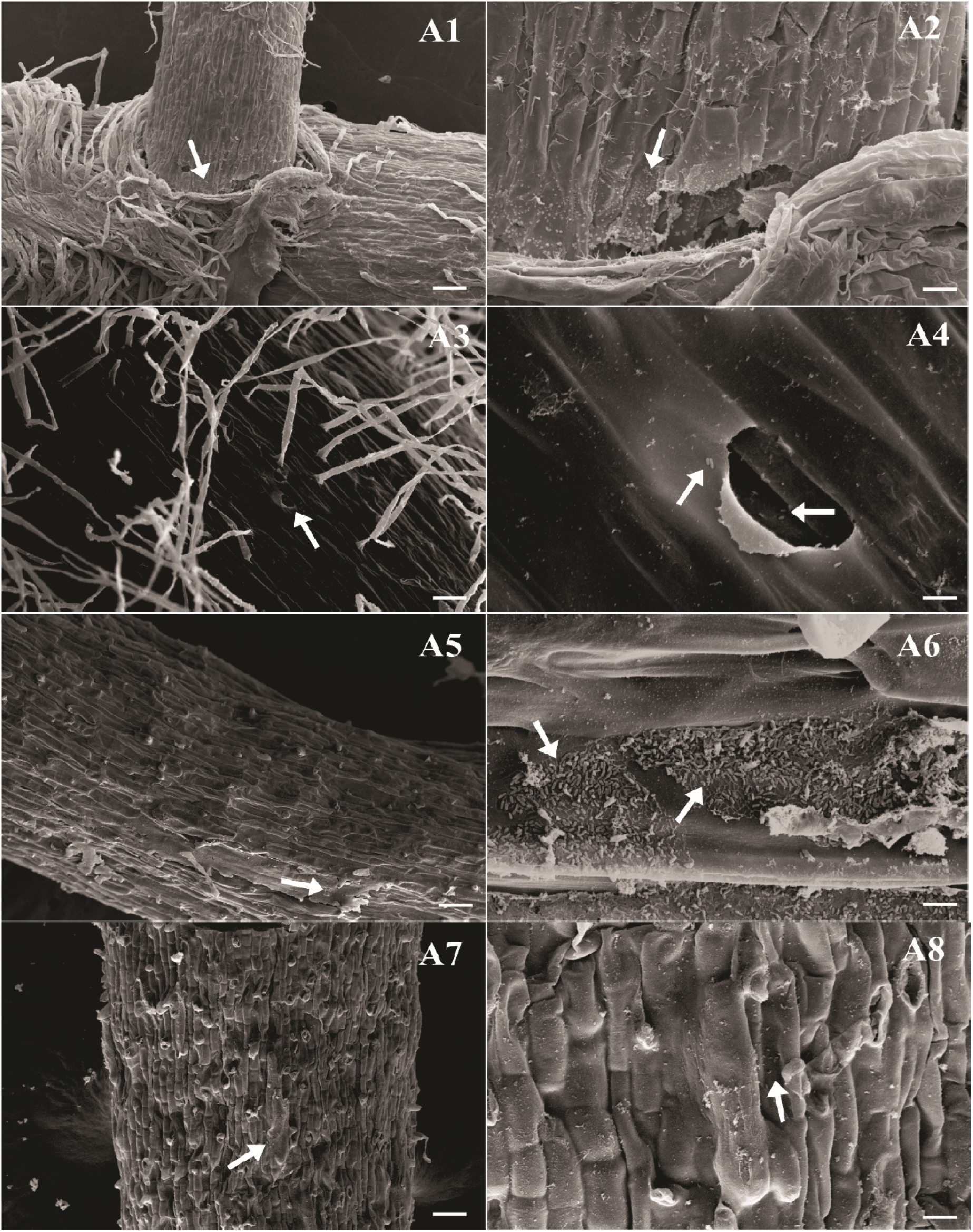
Colonization of disinfected-inoculated maize roots (A). Bacterial cells were visualized by scanning electron microscopy. Root regions: lateral root (A1 and A2), root hair (A3 and A4), growth region (A5 and A6), and cap (A7 and A8). White arrows indicate biofilms and small bacterial aggregates. Bars represent the following scales: panel A4 and A6: 10 μm; A2 and A8: 20 μm; A3, A5, and A7: 100 μm; A1: 200 μm.

### Biochemical analysis of seed reserves under seedling germination

All seeds were treated and germinated according to item “2.1”, with some modifications – including immersion of the non-disinfected group in sterile water for 35 min, while the second group was disinfected in ethanol/hypochlorite for the same time. Stored seed reserves were quantified from three compartments of maize: 1) whole seed: sampled after the soaking and disinfection phase; 2) embryonic axis: collected 24 (for NDS) and 48 h (for DS and DS + MIX) after radicle protrusion; 3) seedling root: obtained after five days of germination (Figure 11). The collected material was macerated in liquid nitrogen and analyzed for protein, glucose, triglyceride, reducing sugar, and alpha-amylase activity. Three replicates per treatment were used. For protein dosing, the bicinchoninic acid (BCA) method (Smith *et al*., 1985) was used, characterized by the reduction of copper ions (Cu^+2^ → Cu^+1^) and formation of the violet BCA-Cu^+1^ complex. The protein concentration was measured at 562 nm in a spectrophotometer. The albumin curve was used as a standard. Reducing sugars were quantified using the 3,5-dinitrosalicylic acid (DNS) method (Miller, 1959). In this method, the sugars react with the DNS (yellow), which is reduced to 3-amino-5-nitrosalicylic acid (dark red). Absorbance was measured at 540 nm. Colorimetric methods determined glucose, triglyceride levels, and alpha-amylase activity and according to the manufacturer’s instructions (Bioclin® K082, K117, K003). Hydrogen peroxide in the presence of specific reagents forms cherry-colored and cherry-red compounds, whose intensity (500-505 nm) is proportional to the concentration of triglyceride and glucose, respectively. The alpha-amylase activity in the samples was inversely proportional to the intensity of the blue color, a product of the complexation between iodine and non-hydrolyzed starch, and calculated by comparing it to a control substrate (600 nm).

**Fig 11.**
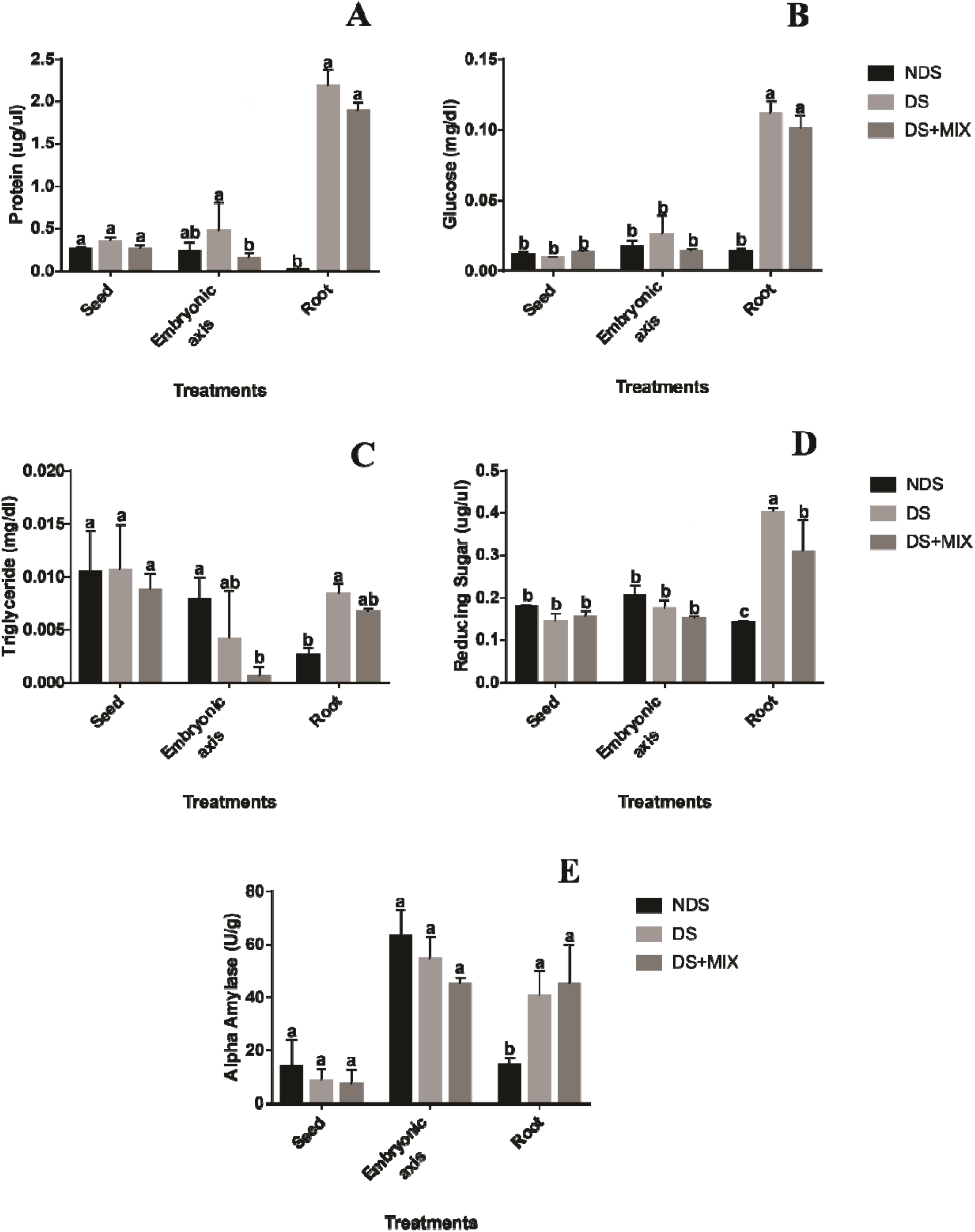
Dosage of protein (A), glucose (B), triglyceride (C), reducing sugar (D), and alpha-amylase activity (E) in different compartments of maize; disinfected or not. *Significant difference between treatments according to the Tukey test (p ≤ 0.05).

## RESULTS

Disinfection assay showed that the increase of the immersion time with sodium hypochlorite (1.25%) proportionally reduced the bacteria population associated with seeds (Fig S1). The partial removal of the microbiota was confirmed by bacterial counts in culture media with different carbon sources. Using LGI and JNFb semi-solid medium, diazotrophic bacteria were not recovered from roots in germinated disinfected seeds. For non-disinfected seeds, values greater than 10^7^ bacteria cells. g^−1^ of fresh root were recovered (Table 2). The isolates obtained as potential diazotrophs had grown in semi-solid medium without adding nitrogen and formed a veil-like pellicle in the subsurface of the medium. In the NB and NB “overnight” medium (NB liquid medium-enriched overnight and plated on solid NB) the total bacterial recovered from disinfected seeds was lower (3.4 × 10^7^ and 5.6 × 10^7^, respectively), while values above 10^8^ were observed in non-disinfected seeds (8.3 × 10^8^ and 4.3 × 10^8^, respectively) (Table 2).

**Table 2.** Differential abundance of roots from disinfected *versus* non-disinfected seeds.

| **rab.all** | **rab.win.R_**<br>**Root_DS** | **rab.win.R_**<br>**Root_NDS** | **effect** | **overlap** | **we.ep** | **Family** | **Genus** |
| --- | --- | --- | --- | --- | --- | --- | --- |
| 14.3029290 | 8.2025510 | 16.6679985 | 3.6336860 | 0.0003653998 | 0.0008595304 | f_Burkholderiaceae | *g_Burkholderia* |
| 1.0463097 | 5.3631248 | -0.4587697 | -2.1410903 | 0.0052604506 | 0.0065642852 | f_Oxalobacteraceae | NA* |
| 4.2163306 | 7.3515686 | 3.0277565 | -1.9246204 | 0.0003653998 | 0.0092850702 | f_Alcaligenaceae | NA* |
| 6.6462458 | 2.1751503 | 8.8878914 | 2.4701196 | 0.0003653998 | 0.0215379059 | f_Burkholderiaceae | *g_Burkholderia* |
| 2.2337604 | 4.2573698 | 0.5699101 | -1.4411406 | 0.0518185718 | 0.0271257626 | f_Corynebacteriaceae | *g_Corynebacterium* |
| 8.2251787 | 2.1263053 | 10.6305670 | 2.6413734 | 0.0003653998 | 0.0274517615 | f_Burkholderiaceae | *g_Burkholderia* |
| 7.3891091 | 1.7284770 | 9.3867903 | 2.7051190 | 0.0003653998 | 0.0297697949 | f_Burkholderiaceae | *g_Burkholderia* |
| 3.5135809 | 5.1803706 | 2.8285070 | -1.8998283 | 0.0259172269 | 0.0299101109 | f_Staphylococcaceae | *g_Staphylococcus* |
| 8.1816592 | 2.1437384 | 10.3636854 | 2.6321218 | 0.0003653998 | 0.0353067527 | f_Burkholderiaceae | *g_Burkholderia* |
| 6.1747742 | 0.4890293 | 6.5897098 | 1.8652017 | 0.0104432569 | 0.0358160707 | f_Burkholderiaceae | *g_Burkholderia* |
| 0.9485628 | 3.6297661 | -0.6027736 | -1.2680380 | 0.0466378252 | 0.0382945172 | f_Microbacteriaceae | *g_Cryocola* |
| 5.7504276 | 0.5893332 | 6.3364622 | 1.7958206 | 0.0207385644 | 0.0456794252 | f_Burkholderiaceae | *g_Burkholderia* |
| 6.1834278 | 0.2673052 | 6.6261919 | 1.8131076 | 0.0103891482 | 0.0515493973 | f_Burkholderiaceae | *g_Burkholderia* |
| 2.7629464 | 4.3992226 | 1.2307604 | -0.8633717 | 0.0937527019 | 0.0536003332 | f_Sinobacteraceae | NA* |

<sup>a</sup>rab.all - median clr value for all samples in the feature
<sup>b</sup>rab.win.No - median clr value for the group of non-disinfected samples
<sup>c</sup>rab.win.Yes - median clr value for the group of disinfected samples
<sup>d</sup>we.ep - Expected P-value of Welch's t-test
\*NA = unassigned

Up to one hundred purified colonies were obtained in the present study (Figure S2A). Analysis of bacterial diversity based on colony morphotype showed that they were more abundant in non-disinfected roots for both culture mediums analyzed (Figure S2B and S2D). The Shannon index, which estimates species richness, was higher in disinfected seed roots (Figure S2C and S2E). Only two types of colonies were shared between treatments (Figure S2F).

Structural analysis of disinfected and non-disinfected maize seeds after imbibition and four days after germination were carried out, aiming to characterize seed-borne microbiota. The surface of the pericarp external layer of non-germinated seeds was colonized by bacteria aggregates in monolayer pattern (Fig S3A1 and S3B1). However, bacteria populations associated with non-disinfected seeds were more abundant. Cross-section of maize kernel also revealed the presence of bacteria in inner tissues colonizing the region of the endosperm (Fig S3A2 and S3B2) and embryo (Fig S3A3 and S3B3) with no apparent differences in bacteria density. With the rupture of the integument by the primary root, more bacteria were visualized by SEM in both treatments. The bacterial aggregates were mainly localized between the pedicel (tip cap) and bottom of the emerged radicle (Fig S3B4) as well in the root hair formation zone of the radicle. Disinfected seeds remove epiphytic bacteria from the pericarp, but endophytic colonization remains unchanged.

The effect of seed-disinfection on the colonization pattern of the microbiota of root was confirmed by light microscopy coupled with TTC-staining (Figure 1) and scanning electron microscopy (Figure 2). It was observed in Figures 1A and 2A of emerged roots from non-disinfected seeds after five days of growth in axenic conditions extensive colonization by the microbiota, mainly formed by bacteria shape cells. In this case, bacterial biofilms were seen more frequently, mainly in lateral roots emergence sites (Fig 2A1-2A2), the root hair zone (Fig 2A3-2A4), and root cap (Fig 2A5-2A6). Microbial colonization in roots of disinfected seed had shown the same colonization sites described for non-disinfected seeds, however, with reduced established population, where single cells and small aggregates were seen more frequently (Fig 1B and 2B). The bacteria cell morphology diversity of the bacterial microbiota was also affected by the disinfection of the seeds, which removed small bacteria cells, in ovoid or short rod shape, and selected long rod shape (Fig 2A1 and 2B4). Interestingly, fungus hyphae were frequently seen in roots that emerged from disinfected seeds (data not shown).

Seed microbiome studies were carried out to understand the seed-borne bacteria community in natural and disinfected seeds of maize variety SHS 5050. First, we analyzed treated and non-germinated seeds. Illumina MiSeq sequencing of the seed bacteriome produced a total of 774 reads for six samples and an average sample coverage of 58% (Supplementary Table S3). Principal coordinate analysis (PCoA) based on the Bray-Curtis dissimilarity matrix suggests that disinfection shapes the seed bacteriome structure and promotes homogeneity within the treatment. Non-disinfected seeds have shown a more heterogeneous pattern. The alpha diversity (within the sample) of the seed bacteriome, measured by the Shannon index, did not differ between the disinfection treatments (p = 0.77) (Supplementary Figure S4). Permanova results showed that seed disinfection did not significantly influence the structure of its bacterial community in no germinated seeds (R2 = 0.178; p = 0.8) (Supplementary Table S4). Venn diagrams were used to show the shared and unique OTUs in each treatment. Seeds treated or not with hypochlorite shared 2 OTUs. The number of unique OTUs was higher in disinfected seeds and not detected in seeds with no disinfection (Supplementary Figure S5). The main bacterial taxa of the seed were classified at the genus level, except for those not assigned, being classified at the family level. f_Enterobacteriaceae_922761 (unassigned genus) were found in disinfected seeds with high relative abundance, composing their main diversity, which was not observed in non-disinfected seeds. The other bacteria indicated in the heatmap (Figure S5) were identified in both treatments but detected in low proportions in the seed bacteriome. Most of the bacteria observed belong to the phylum Proteobacteria (7 out of 10 genera), followed by Actinobacteria and Firmicutes in equal proportion (2 out of 10 genera).

Also, the bacteria community recovered from roots of disinfected and non-disinfected maize seeds germinated under gnotobiotic conditions was evaluate. We sequenced 16S amplicons from 12 maize root samples, which produced a total of 250,303 reads and 88% average sample coverage (Supplementary Table S5).

Ordering by PCoA representing beta diversity shows that disinfection with hypochlorite shapes the maize root bacteriome (Figure 3). The structure of the bacterial community differs between roots from disinfected and non-disinfected seeds. Alpha diversity measures indicated significant differences between the roots analyzed (p = 0.0051), with greater diversity in the treatment whose bacteriome was removed by superficial disinfection of the seed (Figure 3). Ordering results were confirmed by Permanova, where the disinfection factor was significant (p = 0.006), contributing 36% of the variation (Supplementary Table S6).

Venn diagram shows a higher number of unique OTUs in roots with unchanged bacteriome (51), followed by a lower number in the disinfected treatment (5). The intersection between treatments shared 4 OTUs (Figure 4). The relative abundance of bacteria in disinfected versus non-disinfected roots, five genera were abundant in each treatment (Fig. 4). In the roots whose bacteriome was not removed, the genus *Burkholderia* was the most abundant, followed by f_Enterobacteriaceae_922761 (unassigned genus), *Rubrobacter*, *Lactobacillus*, and *Azospirillum*. In the disinfected treatment, the most abundant bacteria belong to the genus *Pseudomonas*, *Bacillus*, *Acinetobacter*, f_Alcaligenaceae_575028 (unassigned genus) and *Corynebacterium*. Proteobacteria dominated the maize root bacteriome in both conditions (7 out of 10 genera), followed by the phyla Actinobacteria, Firmicutes, and Bacteroidetes in equal proportion (1 out of 10 genera).

Differential abundance analysis was performed to filter the bacterial genus removed from the maize root by the disinfection process (Table 2). Disinfection with hypochlorite significantly removed (p ≤ 0.05) bacteria of the *Burkholderia* genus, the most abundant group in the maize root microbiome (Fig 4).

Quantification of the seed bacteriome by qPCR showed that disinfection reduced the number of bacterial cells from 5.9 to 3.3 (p ≤ 0.05) (Figure 5A). At the root, although this effect was not significant (NDS: 6.9 log cell; DS: 6.3; p ≥ 0.05) (Figure 5B).

To evaluate the influence of the microbiota on the germination and growth of maize plantlets, disinfected seed with a reduced bacterial population was subjected to a bacterial recomposition using twelve seed-borne isolates obtained in the present study. Members of this synthetic bacteria community were identified by sequencing the 16S rRNA (Table S2). Nine isolates were attributed to the genus *Burkholderia* (*Burkholderia* sp. and *Burkholderia* gladioli); two to the genus *Bacillus* (*Bacillus drentensis* and *Bacillus camelliae*) and one identified as belonging to the genus *Staphylococcus* sp.

The inoculation of the synthetic bacterial community resulted in higher % G and TMG in seeds inoculated with the bacterial mix (82.5%; 2.57 days), when compared to disinfected seeds, without inoculation (52.5%; 3.21 days) (Figure 6). When comparing inoculated seeds (82.5%) and non-disinfected seeds (92.5%), the %G had shown the same behavior with no statistical differences. However, for TMG, the treatments differed (p ≤ 0.05). For IVG and VGM, no significant differences were observed between inoculated seeds and disinfected seeds, only between controls (NDS and DS).

Five days after germination, the length of the root and aerial part of the seedlings from non-disinfected seeds was significantly longer (17 cm; 6.25 cm) than disinfected seedlings (8.62 cm; 2.83 cm), but did not differ from disinfected-inoculated seeds (14.79 cm; 4.95 cm) (Figure 7). Inoculated seedlings showed more significant growth and produced higher amounts of dry root biomass (0.109 g dry matter) than plants treated with hypochlorite, without inoculation (0.063 g dry matter), demonstrating that the microbiota influence germination and plantlets growth. For the other parameters analyzed, no differences were observed between the disinfected and the disinfected-inoculated treatment.

SEM confirmed the high bacterial colonization of disinfected seeds with partial bacteria community restoration. The pericarp cell wall surface was pronouncedly colonized by single cells and aggregates along the cell junctions (Figure 8A1-8A2). In disinfected seeds, epiphytic bacteria (from the pericarp) were removed by the hypochlorite treatment (Figure 8B1). In cross-sectioned inoculated-seeds, we visualized bacteria cells in low density at endosperm and embryo regions (Figure 8A3-8A6). Interesting to quote that the site of radicle emergence was heavily colonized by bacteria aggregates and biofilms (Fig 8A7-8A8). This region seems to be the preferential site for bacterial growth on germinated seeds since seed-borne bacteria naturally colonized it in non-disinfected seeds, as previously showed in this study.

Light microscopy combined with TTC staining and SEM disinfected-inoculated roots densely colonized by bacteria (Figures 9-10) compared with disinfected treatment (Figures 1B-2B). Massive colonization was observed in regions with lateral root emission and root cap (Fig. 9). More significant bacterial aggregation was identified at points of emergence of lateral root and growing zone. In the root-hair zone and root cap, the cells were more dispersed (Fig. 10). The intense bacterial colonization in the disinfected-inoculated treatment was confirmed by colony count, where the number of CFU in inoculated roots was 1.3 × 1010, compared to 4.5 × 10^7^ in roots treated with hypochlorite and 2.1 × 10^9^ in non-disinfected treatment.

Analysis of the stored reserve mobilization in germinated seeds was accessed in three seed compartments (roots, embryonic axis, and seed, Figure 11). Data from roots showed that glucose degradation was reduced in seeds treated with hypochlorite and reinoculated.

Non-disinfected treatment, the glucose was quickly driven to respiration or cell wall formation. There was also less protein degradation in maize roots that had their microbiota recomposed. In the embryonic axis compartment, the disinfected seeds showed low degradation of their total protein reserve, differing from the other treatments (Figure 11A-B). The degradation of stored triglycerides was more considerable in the non-disinfected roots and the reinoculated embryonic axis and remained unchanged in the seed compartment. A more significant breakdown of triglycerides was observed, which did not occur for glucose and protein (Figure 11C). In the maize root, reducing sugars showed a significant increase in the disinfected treatment, followed by the reinoculated treatment. The rapid mobilization of reserves in non-disinfected seeds resulted in a low content of this sugar. The treated seeds did not differ in terms of reducing sugar content in the region of the embryonic and root axis (Figure 11D). The activity of the alpha-amylase enzyme did not change between treatments in the seed compartment. In the region of the embryonic axis, the enzyme reached higher activity with 24/48 h of germination, with no difference between treatments. Removing part of the maize microbiota increased the activity of alpha-amylase in the roots. The same was observed for the reinoculated roots (Figure 11E).

## DISCUSSION

The present work unravels the bacterial community structure from seed to maize-root seedlings under the gnotobiotic system. Seed-disinfection with sodium hypochlorite was performed, aiming to reduce the natural seed microbial inhabitants to access the functional roles of the seed-microbiome. The comparative analysis emphasizes the importance of the bacterial microbiome to seedlings growth rates and stored reserve mobilization.

The multiple events of germination begin after water is absorbed by the seed and is completed with the radicle emission, a structure that will give rise to a differentiated root (Bewley 1997: 1055). During this process, not only the seeds but also its microbiome, change from a quiescent state to an active physiological state. According to (Frank, et al. 2017), the seeds exude carbon in the form of sugars, proteins, and fatty acids, which act as an energy source for microorganisms and have the potential to shape the bacterial composition around the seed. In response to the availability of nutrients released from the seed or water, microbial cells leave their dormant state and are transferred to seedlings and various organs of mature plants (Nelson, et al. 2018: 1-5), where they can have direct consequences on their germination and growth.

The contribution of the microbiome to plant physiology is still not entirely understood. Herein, we demonstrated that the partial removal of seed-borne bacteria by rigorous disinfection of maize seeds in sodium hypochlorite significantly reduced seed germination speed and seedling growth. This result can be justified by the hypochlorite’s antimicrobial action, which killed relevant bacteria groups modulating germination and growth processes. Increased seed-disinfestation time reduces bacteria population communities and eliminates certain species lowering seedlings germination. Similar results were found by (Verma and White 2018: 764-78, Verma, et al. 2018: 223-38) when increasing the sterilization time in NaOCl from 20 to 40 min. In similar studies, seedling growth was also restricted by removing epiphytic and endophytic microorganisms from seeds through superficial sterilization with NaClO, use of antibiotics (Verma and White 2018: 764-78, Verma, et al. 2018: 223-38, Verma, et al. 2017: 1680-91) or heat treatment (Holland 2016: 31-42, Holland 2019: 21-34).

Our data revealed that the disinfection of seeds with hypochlorite (even for a long time as 90 minutes) does not guarantee the complete elimination of microorganisms, especially those living inside the seed. In roots emerged from untreated-seeds, bacterial cells were viewed in the microscope as aggregates and biofilms as observed by LM and SEM techniques. Since the cultivation system was axenic, the bacteria community described come from the seed itself and can proliferate during seed germination and seedling development. Thus, it is claimed that part of the initial bacterial community of maize seedlings, detected by different microscopies, originates from the seed itself, by vertical transfer (Frank, et al. 2017, Johnston-Monje, et al. 2016: 337-55).

For disinfected seeds, only surface-living resistant microbes or inhabitant of inner tissues were able to colonize emerged primary roots. In this case, the colonization pattern was affected, with reduced bacteria density, mainly viewed as single cells or small groups with cell morphology distinct from the control (natural seed inhabitants). Surface disinfection methods usually carried out with ethanol and sodium hypochlorite, eliminate, or inactivate mainly epiphytic microorganisms from the seed, but not those occupying endophytic sites.

The antimicrobial effect of the hypochlorite was demonstrated by the absence of diazotrophic bacteria and a reduced number of total bacteria associated with emerged roots from disinfected seeds. In control, due to the absence of hypochlorite, the bacterial count was high for all the culture media used. The partial removal of the root bacteriome associated to the disinfested seeds was confirmed by qPCR. At the root, the reduction of bacterial cells in the disinfested treatment was not significant. The high quantification of bacteria at the root of the disinfected treatment can be explained by the niche theory, which predicts that the microbial diversity of plants is determined by the number of available niches (Hardoim, 2015). In this study, the disinfestation of seeds “vacated” niches in the corn root, which started to be colonized by other bacteria (endophytic or resistant to hypochlorite).

Cultivation-dependent approach and cultivation-independent sequencing by Illumina MiSeq revealed similar trends into the bacteriome shift for treated-seeds in comparison with natural seeds. For cultivable fraction, non-disinfected seeds had their colony diversity (richness) reduced during the emergence of the roots, with the predominance of few colony morphotypes (abundance of specific groups). Contrary to this, disinfected seeds displayed the increased richness of colony morphotypes and decreased abundance of specific colonies morphotypes. Similar results were obtained in root sequencing analyses, which exhibited high bacterial diversity by the Shannon index when the hypochlorite removed part of its bacteriome. Diversity values were lower in non-disinfected seeds. These results are justified by the increase in the abundance of common or predominant taxa in untreated corn seed, perhaps those from epiphytic sites, and the elimination of transient microorganisms. In the disinfected treatment, data reinforce the selection of colonizers resistant to sodium hypochlorite or endophytic bacteria resident from embryo or endosperm of the seed. With the removal of bacteria prevalent in the seed, other taxa could be detected by the sequencing technique.

Sequencing results of the 16S rRNA gene showed a low number of reads associated with no germinated seeds for disinfected and non-disinfected treatments. Additionally, hypochlorite treatment did not significantly affect bacteria diversity of the seed-borne community. However, the abundance of bacteria from the Enterobacteriaceae family was evident in the seeds treated with hypochlorite. Bacteria in this group often occupy the deepest tissues of plants (Liu, et al. 2017: 317-24), which justifies their detection in this treatment. Besides, members of the Enterobacteriaceae family may be resistant to antimicrobial compounds (Wellington, et al. 2013: 155-65) or rapidly consume the seed exudates, which explains their growth under these conditions (Nelson 2004: 271-309). From the seed, the Enterobacteriaceae family colonized, to a lesser extent, the corn root, suggesting that competition with other bacteria may have reduced its dominance (Johnston-Monje, et al. 2016: 337-55).

For bacteria groups associated with emerged radicles, the genus *Burkholderia* was selectively removed by the hypochlorite treatment. In contrast, roots that showed higher growth rates (non-disinfected treatment) were abundantly colonized by *Burkholderia*. This genus was defined by (Yabuuchi, et al. 1992: 1251-75) and occupied diverse ecological niches (Coenye and Vandamme 2003: 719-29). *Burkholderia* species can colonize diverse plant host, and thus promote their growth and protection against pathogens (Tagele, et al. 2018: 8-18, Tagele, et al. 2019: 1005). *Burkholderia* abundance has been reported in maize and other plants (Johnston-Monje, et al. 2016: 337-55, Mashiane, et al. 2017: 80, Rosenblueth, et al. 2010: 39-48), been able to fix nitrogen, solubilizing phosphate, producing AIA (indole-acetic acid; auxin) and siderophores (Batista, et al. 2018: 33-42). Genomic studies had shown several genes related to the biosynthesis of AIA, ACC deaminase, dioxygenases, multiple efflux pumps, degradation of aromatic compounds, and diverse protein secretion systems (Dias, et al. 2019: e00801). Some of these genetic traits may have contributed to the growth of seedlings with *Burkholderia* in its bacteriome.

Twelve bacteria isolated from the non-disinfected seeds were used to compose a synthetic bacteria community aiming to evaluate the microbiota recomposition phenotype. Nine from twelve bacterial isolates obtained were identified as belonging to the *Burkholderia* genus. After inoculation in disinfected seeds, the germination and growth of the maize seedlings were recovered, being comparable to the development rates of seeds that were not treated with hypochlorite. Bacteria counts and microscopy analysis confirmed the partial rebuilding of the microbiota. In similar studies, it has been found that removing the microbiota from seeds has reduced the germination and development of rice, soybeans, beans, and millet (Verma and White 2018: 764-78, Verma, et al. 2018: 223-38). (Holland 2019: 21-34) used thermotherapy at 50 °C for 48 h for seed disinfection on soybean seeds and demonstrated a population reduction of *Methylobacterium* by 3% associated with reduced germination and abnormal seedling growth. All of the growing and developmental plant traits were restored after the reintroduction of removed bacterial community part. Collectively, these studies suggest that the germination and growth rates of seedlings can be positively modulated, at least in part, by the bacteriome associated with seeds. The increased seedling fitness can be related to the production of phytohormones, synthesis of other metabolites and alleviation of damage caused by pathogens, through the production of antifungal and antibacterial compounds.

The plantlets’ growth performance can be partially associated with the bacteriome. The underlined mechanism can be, in part, related to the positive modulation of seed reserve mobilization during germination. Thus, for root from non-disinfected maize seeds, protein, glucose, and triglyceride reserve degradation is more efficient than when the associated microbiota was partially removed. The low content of reducing sugars and the increase of the activity of the alpha-amylase enzyme in the embryonic axis supports rapid plant growth in intact and restored microbiota. On the other hand, the disinfection of the seeds removed microbiota bacteria and promoted late mobilization of its biomolecule reserves, evidenced by the accumulation of protein, glucose, triglyceride, and reducing sugar in the root, in addition to high alpha-amylase activity in the post-germination phase. Late mobilization of reserves affected maize growth. (Zhu, et al. 2017: 2029-35) found similar where Ammodendron bifolium endophytic bacteria increased the efficiency of endosperm use, including more significant degradation of sucrose, protein, and triglyceride from the seed, in addition to the production of hydrolytic enzymes by these bacteria. For (Verma, et al. 2017: 1680-91), seed bacteria can express the enzyme amylase and increase the efficiency of endosperm use during germination and seedling growth. The idea that the microbiota can mobilize nutrients from the endosperm to the embryo is supported by the presence of bacteria in these regions, located by scanning microscopy.

Overall results point that even after the removal of part of seed-borne bacteria, similar population levels were detected by qPCR in roots from disinfected and non-disinfected germinated seeds. Therefore, the remaining bacteriome from disinfected-seeds exhibited selective removal of prevalent fast-grown groups and specific taxon. Then, under less competition, the remaining bacteria can move to the root and colonize it during germination when the seed releases several metabolites that shape the bacterial composition (Frank, et al. 2017). This fact was proven by detecting *Enterobacteriaceae*, *Acinetobacter*, *Pseudomonas*, *Lactobacillus*, and *Burkholderia* in maize seeds and roots by amplicon sequencing. SEM also confirmed the migration of the remaining bacteria from seeds to radicle. However, the population size and diversity of these bacteria in the treatments are distinct. As quoted before, in disinfected treatment, the root bacteriome is more diverse, but this diversity does not include or contemplates a reduced number of bacteria that promote germination and growth, which negatively affected the development of these seedlings.

On the other hand, the natural microbiome of maize roots is composed of more dominant groups and less diverse bacterial taxa, which act positively in the efficient mobilization of seed reserves and processes of biostimulation and biofertilization. It can be inferred from the sequencing results that bacteria of the genus *Burkholderia* play a significant role in these processes since the partial rebuild of the microbiota with members of this genus has recovered maize germination and plant growth processes. Whatever the underlined mechanism displayed, it is evident that physiological processes during germination and seedling growth of maize seeds were modulated by seed-borne bacteriome.

## SUPPLEMENTARY DATA

Table S1. Bacterial isolates obtained from seed and maize root microbiota.

Table S2. Identification of microbiome bacteria by sequencing the 16S rRNA after BLAST in the NCBI database.

Table S3. Distribution of reads per seed sample.

Table S4. Permutational Multivariate Variance Analysis (Permanova) of the seed bacteriome.

Table S5. Distribution of reads by root sample of germinated disinfected and no disinfected seeds variety SHS 5050.

Table S6. Permutational Multivariate Variance Analysis (Permanova) of the root bacteriome.

Fig S1. Effect of disinfection time on bacteria population size and germination parameters associated with maize seeds.

Fig S2. Diversity of bacterial colonies in non-disinfected (NDS) and disinfected (DS) seed roots.

Fig S3. Colonization of non-germinated maize seeds by its microbiota. Bacterial cells were visualized by scanning electron microscopy (SEM).

Fig S4. The principal coordinate graph (PCoA) based on the Bray-Curtis dissimilarity matrix (A) and measurements of the alpha diversity of the bacteriome of disinfected (Yes) and non-disinfected (No) non-geminated seeds.

Fig S5. Venn diagram showing OTUs overlap (A) and the relative abundance between seed from disinfected (Yes) and non-disinfected (No) (B) non-germinated seeds.

## ACKNOWLEDGMENTS

This work is part of the Ph.D. thesis from LFS that acknowledge CAPES for the fellowship.

