## Supplemental Table_Figures for "Insights into the structure and role of seed-borne bacteriome during maize germination"

**Supplementary data**

**Table S1.** Bacterial isolates obtained from seed and maize root microbiota.

**Table S2.** Identification of microbiome bacteria by sequencing the 16S rRNA after BLAST in the NCBI database.

**Table S3.** Distribution of reads per seed sample.

**Table S4.** Permutational Multivariate Variance Analysis (Permanova) of the seed bacteriome.

**Table S5.** Distribution of reads by root sample of germinated disinfected and no disinfected seeds variety SHS 5050.

**Table S6.** Permutational Multivariate Variance Analysis (Permanova) of the root bacteriome.

**Fig S1.** Effect of disinfection time on bacteria population size and germination parameters associated with maize seeds.

**Fig S2.** Diversity of bacterial colonies in non-disinfected (NDS) and disinfected (DS) seed roots.

**Fig S3.** Colonization of non-germinated maize seeds by its microbiota. Bacterial cells were visualized by scanning electron microscopy (SEM).

**Fig S4.** The principal coordinate graph (PCoA) based on the Bray-Curtis dissimilarity matrix (A) and measurements of the alpha diversity of the bacteriome of disinfected (Yes) and non-disinfected (No) non-geminated seeds.

**Fig S5.** Venn diagram showing OTUs overlap (A) and the relative abundance between seed from disinfected (Yes) and non-disinfected (No) (B) non-germinated seeds.

**Table S1.** Bacterial isolates obtained from seed and maize root microbiota.

| **Bacteria strain designation** | **Origin** |
| --- | --- |
| LMS 3b | NB NDS R2 -5 |
| LMS 4 | NB NDS R2 -6 |
| LMS 5 | NB NDS R2 -6 |
| LMS 7 | NB NDS R2 -6 |
| LMS 8  LMS 10 | NB NDS R2 -6  NB NDS R2 -6 |
| LMS 23 | NDS NB overnight R2 -4 |
| LMS 25b | NDS NB overnight R2 -6 |
| LMS 73b1 | LGI NDS R2 -6 (-7) |
| LMS 77  LMS 81  LMS 99 | LGI NDS R4 -5 (-7)  JNFb NDS R1 -4 (-7)  JNFb NDS R4 -6 (-7) |
| Culture media used for isolation: NB solid medium and JNFb and LGI semi-solid medium; Non-disinfected seed (NDS); the number of experimental unit replication and dilution obtained (RX -X). | |

**Table S2.** Identification of microbiome bacteria by sequencing the 16S rRNA after BLAST in the NCBI database.

| **Isolate** | **Isolate identified by 16S rRNA sequence** | **Access number** | **Identity (%)** | **E value** | **Query cover (%)** | **Score** |
| --- | --- | --- | --- | --- | --- | --- |
| 3b | *Staphylococcus sp.* strain Firmi-16 | MH683105.1 | 99.59 | 0.00 | 91 | 1332 |
| 4 | *Burkholderia sp.* strain Beta-30 | MH698880.1 | 99.57 | 0.00 | 100 | 1279 |
| 5 | *Burkholderia gladioli* strain E37CS3 | MK474979.1 | 99.87 | 0.00 | 100 | 1413 |
| 7 | *Burkholderia sp.* strain BIS1062 | MN810235.1 | 99.87 | 0.00 | 100 | 1408 |
| 8 | *Burkholderia gladioli* strain 16BC | MK474977.1 | 100 | 0.00 | 100 | 1232 |
| 10 | *Burkholderia gladioli* strain T3.3.1 | MT001453.1 | 99.85 | 0.00 | 100 | 1247 |
| 23 | *Bacillus drentensis* strain CD3 | MK216756.1 | 100 | 0.00 | 100 | 1330 |
| 25b | *Bacillus camelliae* strain 7578-1 | NR_159341.1 | 99.72 | 0.00 | 98 | 1321 |
| 73b1 | *Burkholderia gladioli* strain E37CS3 | MK474979.1 | 100 | 0.00 | 100 | 1291 |
| 77 | *Burkholderia gladioli* strain TWS (19)-Abf_48Ba | MN049500.1 | 100 | 0.00 | 99 | 1293 |
| 81 | *Burkholderia sp.* strain Beta-30 | MH698880.1 | 100 | 0.00 | 100 | 1291 |
| 99 | *Burkholderia gladioli strain* TWS (19)-Abf_48Ba | MN049500.1 | 100 | 0.00 | 100 | 1293 |

**Table S3.** Distribution of reads per seed sample.

| **Sample ID** | **Disinfection** | **Treatment** | **Total Reads** | **Coverage** |
| --- | --- | --- | --- | --- |
| **O28** | No | S_SHS5050_ND_CTAB | 273 | 0.7728938 |
| **O29** | No | S_SHS5050_ND_CTAB | 56 | 0.4642857 |
| **O27** | Yes | S_SHS5050_D_CTAB | 96 | 0.5416667 |
| **O25** | Yes | S_SHS5050_D_CTAB | 223 | 0.6188341 |
| **O26** | Yes | S_SHS5050_D_CTAB | 84 | 0.5357143 |
| **O30** | No | S_SHS5050_ND_CTAB | 42 | 0.5238095 |

**Table S4.** Permutational Multivariate Variance Analysis (Permanova) of the seed bacteriome.

|  | **Df** | **Sums of Sqs** | **Mean Sqs** | **F. Model** | **R2** | **Pr (>F)** |
| --- | --- | --- | --- | --- | --- | --- |
| **Disinfection** | 1 | 0.308129 | 0.3081290 | 0.8668261 | 0.1781091 | 0.8 |
| **Residuals** | 4 | 1.421872 | 0.3554681 | NA | 0.8218909 | NA |
| **Total** | 5 | 1.730001 | NA | NA | 1.0000000 | NA |

**^a^Df: degrees of freedom**

**^b^Sum of Sqs: sequential sums of squares**

**^c^Mean Sqs: mean squares**

**^d^F. Model: F statistics**

**^e^Pr>(F): partial R-squared and P values**

**Table S5.** Distribution of reads by root sample of germinated disinfected and no disinfected seeds variety SHS 5050.

| **X.Sample**  **ID** | **Genotype** | **Extraction** | **Disinfection** | **Treatment** | **Total Reads** | **Coverage** |
| --- | --- | --- | --- | --- | --- | --- |
| **O12** | SHS5050 | CTAB | No | R_SHS5050_ND_CTAB | 64695 | 0.9944354 |
| **O11** | SHS5050 | CTAB | No | R_SHS5050_ND_CTAB | 23212 | 0.9912545 |
| **O4** | SHS5050 | DNAZol | No | R_SHS5050_ND_DNAZol | 49737 | 0.9941291 |
| **O2** | SHS5050 | DNAZol | Yes | R_SHS5050_D_DNAZol | 1248 | 0.9551282 |
| **O8** | SHS5050 | CTAB | Yes | R_SHS5050_D_CTAB | 118 | 0.6949153 |
| **O9** | SHS5050 | CTAB | Yes | R_SHS5050_D_CTAB | 144 | 0.5555556 |
| **O5** | SHS5050 | DNAZol | No | R_SHS5050_ND_DNAZol | 64020 | 0.9946892 |
| **O3** | SHS5050 | DNAZol | Yes | R_SHS5050_D_DNAZol | 263 | 0.8365019 |
| **O6** | SHS5050 | DNAZol | No | R_SHS5050_ND_DNAZol | 42802 | 0.9939022 |
| **O10** | SHS5050 | CTAB | No | R_SHS5050_ND_CTAB | 3686 | 0.9750407 |
| **O7** | SHS5050 | CTAB | Yes | R_SHS5050_D_CTAB | 148 | 0.7229730 |
| **O1** | SHS5050 | DNAZol | Yes | R_SHS5050_D_DNAZol | 230 | 0.8391304 |

**Table S6.** Permutational Multivariate Variance Analysis (Permanova) of the root bacteriome.

|  | **Df** | **Sums of Sqs** | **Mean Sqs** | **F. Model** | **R2** | **Pr(>F)** |
| --- | --- | --- | --- | --- | --- | --- |
| **Disinfection** | 1 | 1.563833 | 1.5638332 | 5.719781 | 0.3638588 | 0.006 |
| **Residuals** | 10 | 2.734079 | 0.2734079 | NA | 0.6361412 | NA |
| **Total** | 11 | 4.297912 | NA | NA | 1.0000000 | NA |

**^a^Df: degrees of freedom**

**^b^Sum of Sqs: sequential sums of squares**

**^c^Mean Sqs: mean squares**

**^d^F. Model: F statistics**

**^e^Pr>(F): partial R-squared and P values**

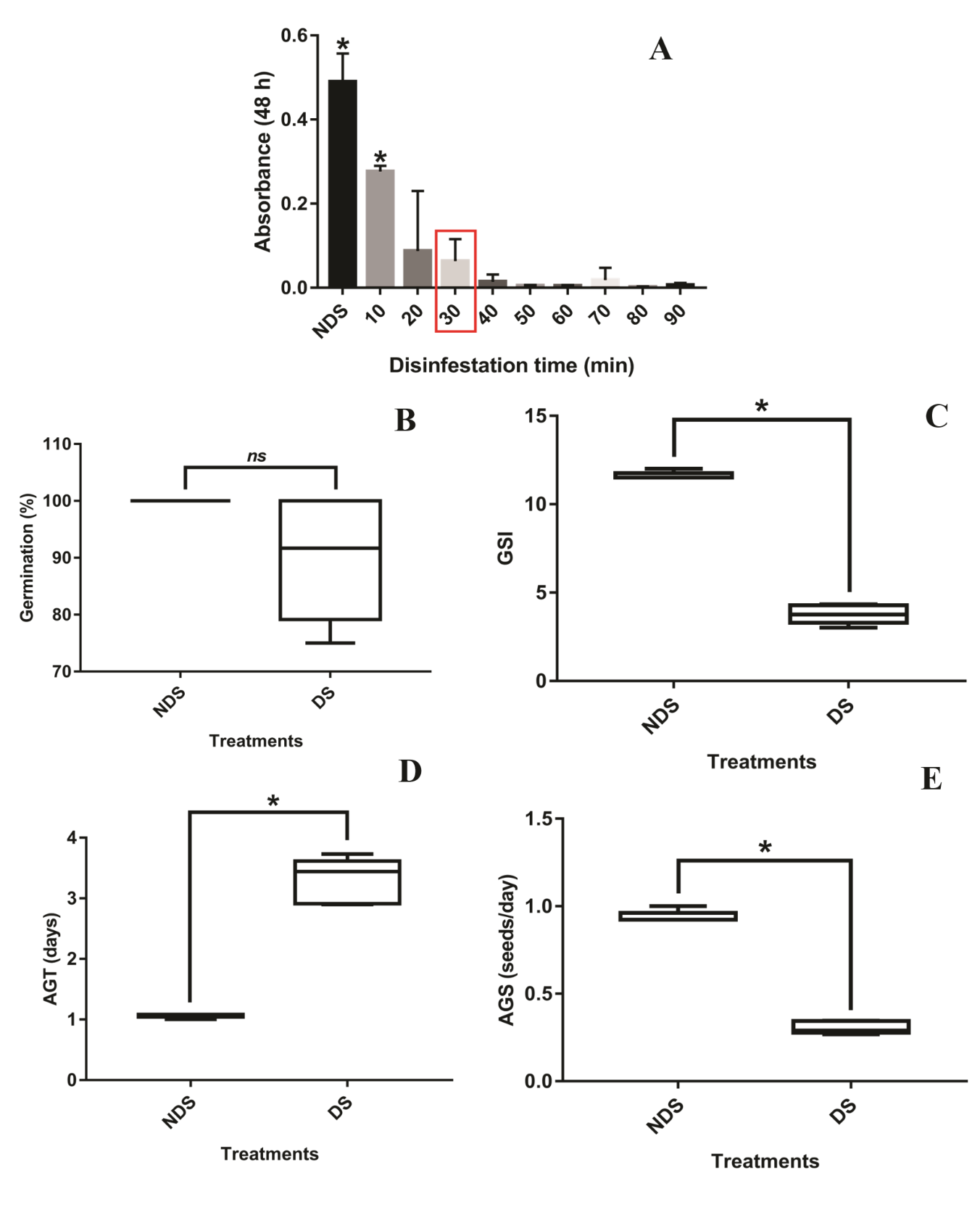

**Fig S1.** Microbial growth reported as an optical density increase in NB liquid medium with maize seeds submitted to different disinfection times in sodium hypochlorite (1.25% for 10, 20, 30, 40, 50, 60, 70, 80 and 90 min) (A). Germination of non-disinfected (NDS) and disinfected (DS) maize seeds (B to E). Germination percentage (B), germination speed index (C), average germination time (D), and average germination speed (E). *Significant difference between treatments according to the Tukey test (p ≤ 0.05).

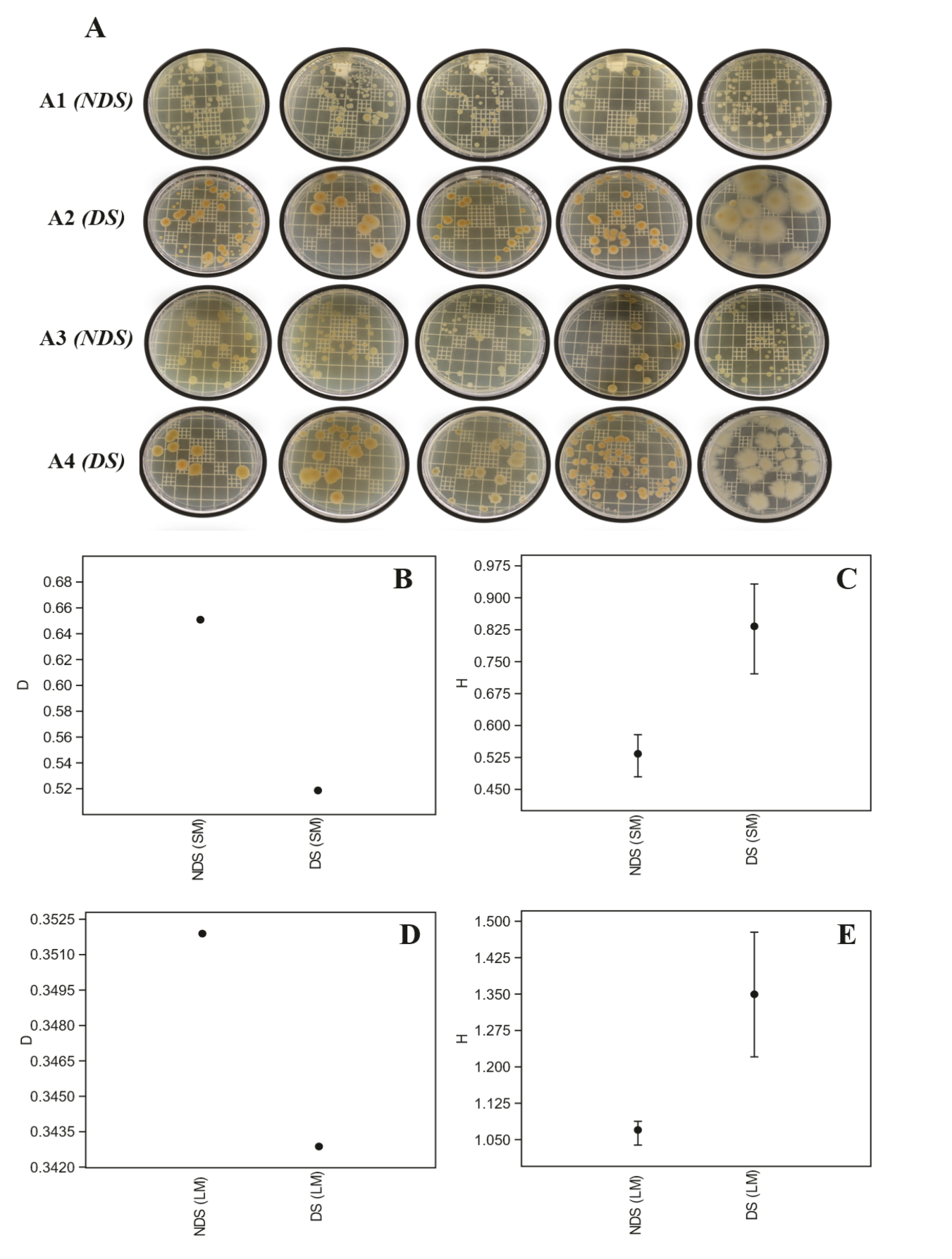

**Fig S2.** Diversity of bacterial colonies in non-disinfected (NDS) and disinfected (DS) seed roots. Bacteria were grown in solid NB medium (SM; A1 and A2) and liquid NB medium “overnight” (LM; A3 and A4), in 5 replicates. Dominance (D; B and D) and Shannon’s Index (H; C and E) were calculated for each treatment.

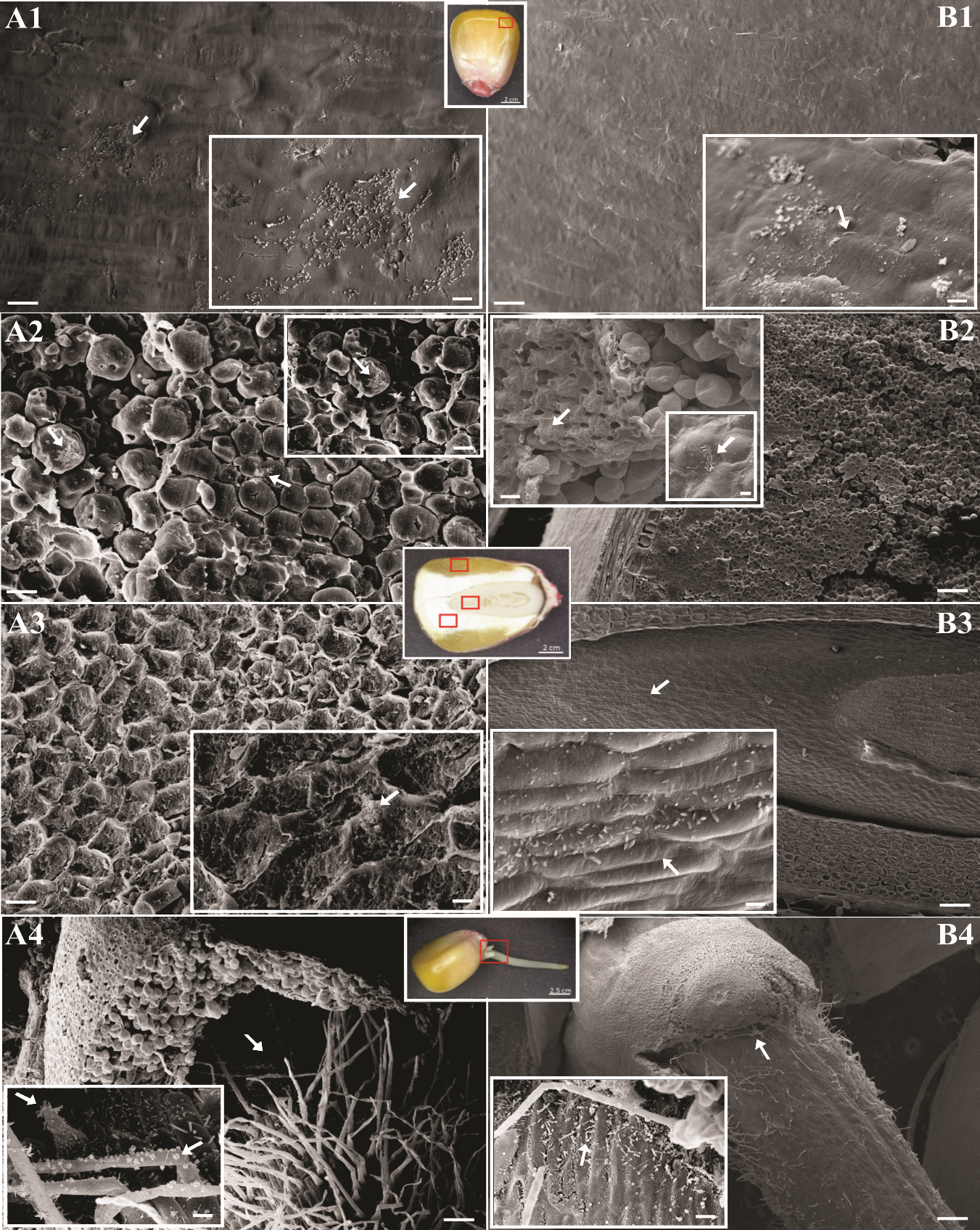

**Fig S3.** Colonization of maize seeds by its microbiota. Bacterial cells were visualized by scanning electron microscopy (SEM). Maize seeds non-disinfected (A) and disinfected (B). Seed regions: pericarp (A1 and B1), endosperm (A2 and B2), embryo (A3 and B3), and radicle (A4 and B4). White arrows indicate biofilms and small bacterial aggregates. Bars represent the following scales: panel A1 and A2: 20 µm; A3 and B1: 20 and 10 µm; A4 and B4: 200 and 20 µm; B2: 100 and 20 µm; B3: 100 and 2 µm.

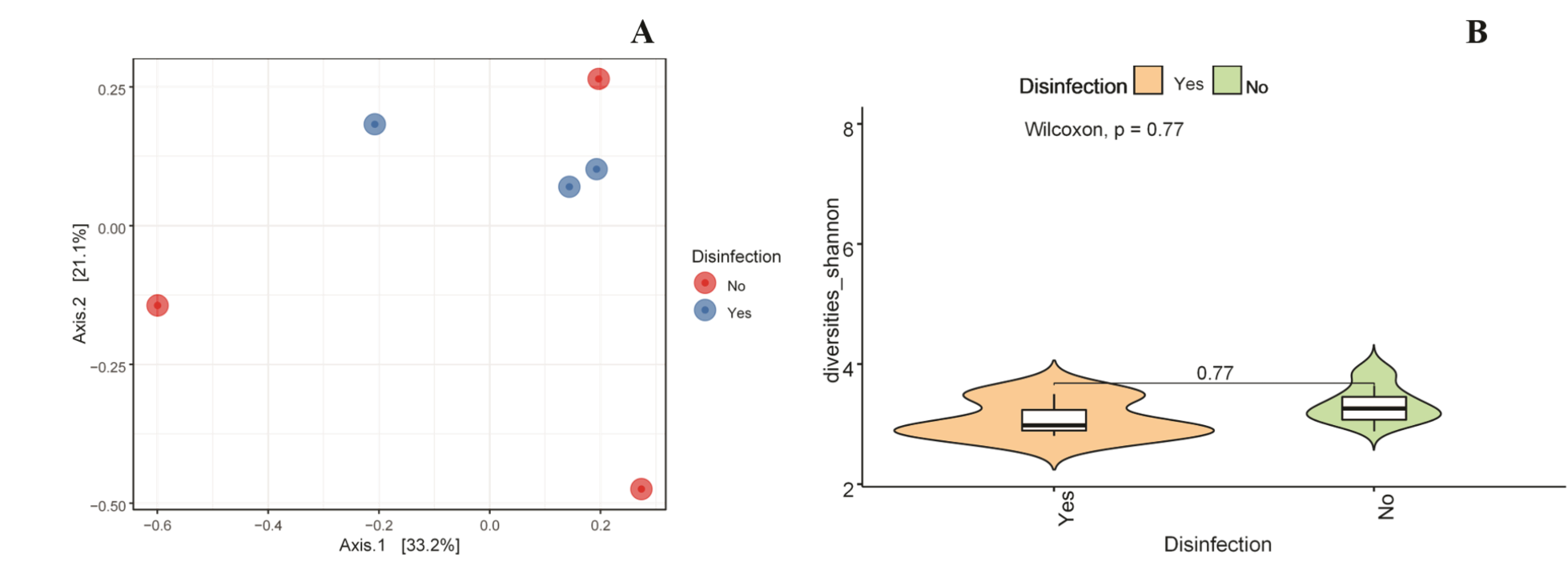

**Fig S4.** The principal coordinate graph (PCoA) based on the Bray-Curtis dissimilarity matrix (A) and measurements of the alpha diversity of the bacteriome of disinfected (Yes) and non-disinfected (No) non-germinate seeds. Different colors indicate the treatments. Shannon = microbial diversity index.

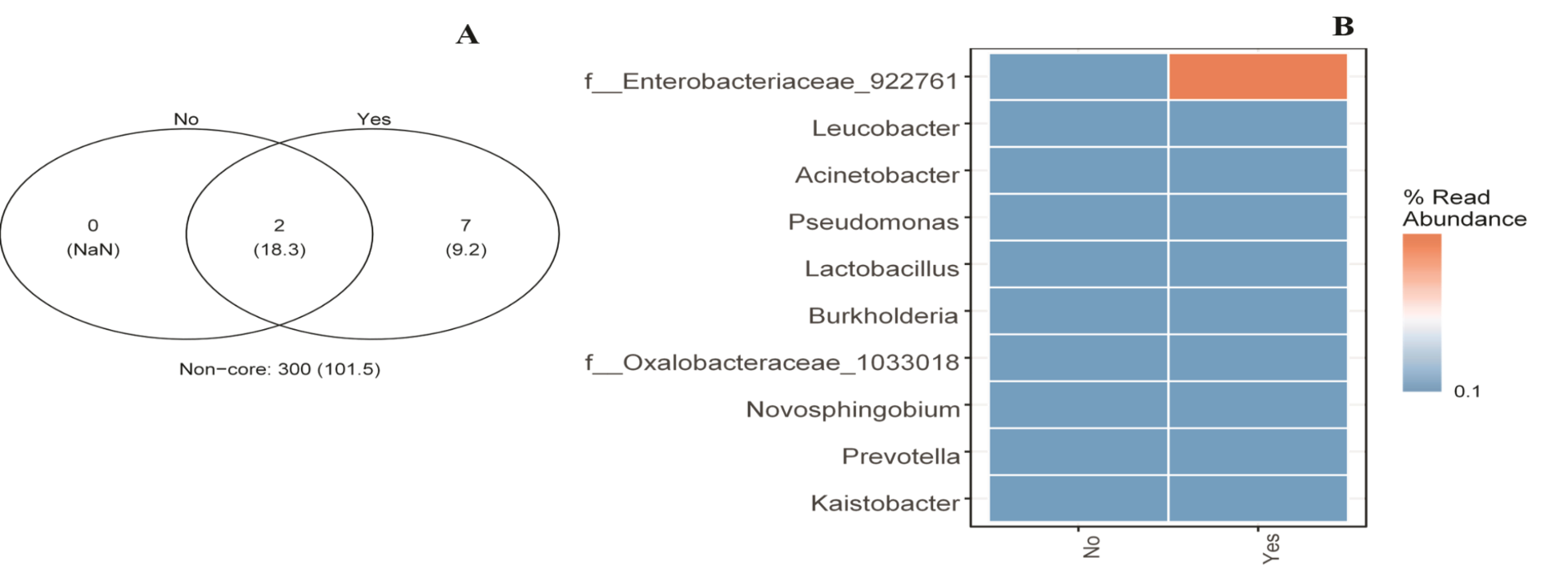

**Fig S5.** Venn diagram showing OTUs overlap (A) and the relative abundance between seed from disinfected (Yes) and non-disinfected (No) non-germinated seeds (B). The color intensity in the heatmap indicated in the legend to the right of the figure shows the relative value.
